# A model of decentralized vision in the sea urchin *Diadema africanum*

**DOI:** 10.1101/2022.05.03.490537

**Authors:** Tianshu Li, John Kirwan, Maria Ina Arnone, Dan-Eric Nilsson, Giancarlo La Camera

**Affiliations:** Department of Neurobiology & Behavior, Stony Brook University, Stony Brook, NY, USA; Program in Neuroscience, Stony Brook University, Stony Brook, NY, USA; Center for Neural Circuit Dynamics, Stony Brook University, Stony Brook, NY, USA; Stazione Zoologica Anton Dohrn, Naples, Italy; Lund Vision Group, Department of Biology, Lund University, Lund, Sweden

**Author notes:** Address correspondence to Giancarlo La Camera, Department of Neurobiology and Behavior, Stony Brook University, Stony Brook, NY.

## Abstract

Sea urchins can detect light and move in relation to luminous stimuli despite lacking eyes. They presumably detect light through photoreceptor cells distributed on their body surface. However, there is currently no mechanistic explanation of how these animals can process light to detect visual stimuli and produce oriented movement. Here, we present a model of decentralized vision in echinoderms that includes all known processing stages, from photoreceptor cells to radial nerve neurons to neurons contained in the oral nerve ring encircling the mouth of the animals. In the model, light stimuli captured by photoreceptor cells produce neural activity in the radial nerve neurons. In turn, neural activity in the radial nerves is integrated in the oral nerve ring to produce a profile of neural activity reaching spatially across several ambulacra. This neural activity is read out to produce a model of movement. The model captures previously published data on the behavior of sea urchin *Diadema africanum* probed with a variety of physical stimuli. The specific pattern of neural connections used in the model makes testable predictions on the properties of single neurons and aggregate neural behavior in *Diadema africanum* and other echinoderms, offering a potential understanding of the mechanism of visual orientation in these animals.

## 1 Introduction

Sea urchins are a large clade of echinoderms found at all depths in the ocean with diverse feeding and behavioral habits. Despite lacking eyes, these marine animals can visually resolve objects and move towards them, as well as point their spines towards looming visual stimuli (Holmes, 1912; Millott and Takahashi, 1963; Yoshida, 1966; Blevins and Johnsen, 2004; Yerramilli and Johnsen, 2010; Al-Wahaibi and Claereboudt, 2017; Kirwan et al., 2018). The long-spined sea urchin *Diadema africanum* (Rodríguez et al., 2013), a night-active, herbivorous diadematoid of the eastern Atlantic which inhabits shallow seas and displays relocation behavior irrespective of the time of day, moves towards dark objects with a spatial resolution of 29–69°, i.e., 0.01-0.03 cycles per degree (Kirwan et al., 2018). Such behavior is normally associated with a centralized brain processing visual information from discrete eyes, yet sea urchins lack both. Therefore, sea urchins possess a unique visual system which has not been studied in terms of its information processing. Here, we set out to provide a mechanistic model of directional, low resolution vision in sea urchins building on our knowledge of their general neural anatomy and the behavior observed in *D. africanum*. By providing such a model we also aim to address a growing interest in the study of neural processing mechanisms in decentralized nervous systems (see e.g. (Clark et al., 2019; Kirwan et al., 2018; Al-Wahaibi and Claereboudt, 2017; Beer et al., 2016; Li et al., 2015; Garm and Nilsson, 2014)). We especially aim to complement recent studies on visual resolution and locomotion in echinoderms with a quantitative framework of how sensory input integration and visual detection can be achieved by their decentralized systems.

In adult sea urchins, resolving vision may be mediated by photoreceptor cells (PRCs) situated in the tube feet of the animal. The tube feet emerge from five vertical grooves in the animal endoskeleton called ‘ambulacra’, located with pentaradial symmetry on the body surface. The tube feet are involved in such tasks as locomotion, positioning, cleaning and feeding (Hyman, 1955), but each tube foot also has PRCs (Agca et al., 2011; Lesser et al., 2011; Ullrich-Lüter et al., 2011). The anatomical and molecular description of PRCs in sea urchins come mostly from studies in the species *Strongylocentrotus purpuratus*, where it has been found that PRCs along the tube feet are shielded such that they only detect light from a restricted angle, approximately orthogonal to the body surface (Woodley, 1982; Blevins and Johnsen, 2004; Yerramilli and Johnsen, 2010; Ullrich-Lüter et al., 2011). Due to this property, the PRCs on the tube feet can together provide coarse spatial information as if the entire animal was a compound eye (Blevins and Johnsen, 2004; Ullrich-Lüter et al., 2011). However, sea urchins have no centralized brain which could process light information coming from the PRCs; rather, their internal nervous system comprises five radial nerves (RNs) and one oral nerve ring (ONR). The ONR is a commissure surrounding the mouth and interconnecting the RNs, and therefore is ideally poised to be responsible for both sensory integration and motor coordination, as previously suggested (see e.g. (Yoshida, 1966) and our Discussion).

In this work, we build a theoretical model of the decentralized visual system of these echinoids based on previously published data on the behavior of *D. africanum* in response to light stimuli (Kirwan et al., 2018). In the model presented here, light information from PRCs is processed in the RNs and then relayed to ONR neurons, whose activity is read out to produce visually guided behavior. We apply our model to explain the behavior of *D. africanum* in the presence of isoluminant visual stimuli with a central dark ‘target’ flanked by lighter regions (Kirwan et al., 2018). *D. africanum* also displays a spine-pointing response to looming circular fields subtending 13–25 degrees (Kirwan et al., 2018); here, however, we focus on modeling the movement towards stimuli that are isoluminant to the background as this allows to discriminate spatial vision from phototaxis. In the experiments, the animals were initially placed at the center of a circular arena, with stimuli placed on its outer wall; the animals would then move towards the wall of the arena, and their behavior was analyzed in search for patterns of directional motion. The model presented here makes testable predictions on the behavior in this setup for a large class of visual stimuli, in particular on the three stimuli tested in the experiments of Kirwan et al. (2018). For these stimuli, the distribution of final positions predicted by the model matches, to a remarkable degree, the distribution of final positions reached by the animals.

To our knowledge, this is the first model of decentralized vision and visually-guided behavior in sea urchins. The model combines all known neural processing stages, from PRCs to RN neurons to ONR neurons, including a probabilistic readout mechanism of the activity of ONR neurons that is responsible for visually-guided movement. The specific pattern of neural connections used in the model also makes testable predictions on the properties of single neurons in each processing stage, such as their cell type (inhibitory vs. excitatory), their connectivity structure, and their response properties. All of these properties can in principle be measured in experiment and may lead to an understanding of the mechanism of visually guided orientation in echinoderms.

## 2 Results

### 2.1 Model of decentralized vision in sea urchins

#### General anatomical features

We constructed a model of vision in *Diadema africanum* by integrating three main components: photoreceptor cells (PRCs), radial nerves (RNs) and oral nerve ring (ONR). PRCs and RNs were located along each of the five ambulacra; the PRCs were located outside the test, along tube feet (Ullrich-Lüter et al., 2011), while the neurons were located inside the test along the latitudinal direction (along the *θ* angle in Fig. 1A). For convenience, we grouped the PRCs in separate groups according to their target neurons in the RNs, and we did the same for the RN neurons (Fig. 1B). The numbers of groups of PRCs and RN neurons in ambulacrum *k* were 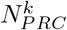 and 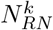, respectively (see Table 1 for model parameter definitions and values). Similarly, we divided the ONR neurons in two subgroups, excitatory and inhibitory, and each subgroup was further divided into *N*_*ONR*_ groups based on the patterns of connectivity (details below). Since sea urchins have five-fold symmetry, the center position of ambulacrum *k* was set to be *ϕ*^*k*^ = 72(*k* −1)°. Since the stimuli were vertically homogeneous (see Sec. “Model prediction of object taxis in *Diadema africanum*”), we simplified the 3D structure of the test to a horizontal slice (along the longitudinal angle *ϕ*, see Fig. 1A).

**Figure 1:**
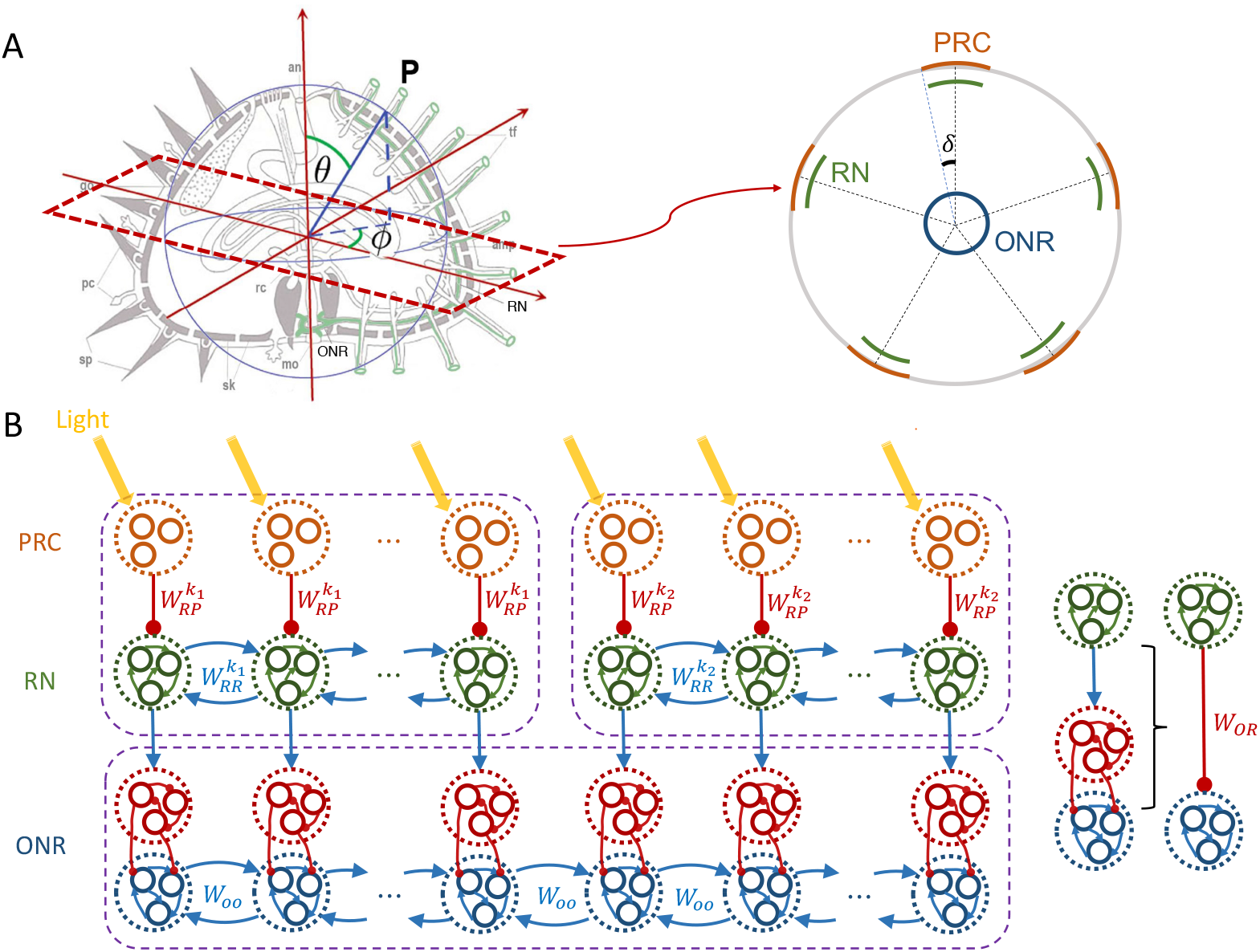
Schematic illustration of the model nervous system. **A** Schematic geometry of the animal. Left: 3D geometry. *θ*: latitudinal angle; *ϕ*: longitudinal angle. *Right:* transverse view of the animal cut above the ONR. The orange arcs indicate the distributions of PRCs on each ambulacrum (*δ*: half width of the distribution). The green arcs indicate RN cells. The blue circle indicates eONR cells. **B** Cartoon of the network structure used in the model (only 2 ambulacra shown at the top, represented by purple rectangles). Red segments terminating in a circle indicate inhibitory connections, blue arrows indicate excitatory connections. From top to bottom, orange circles indicate groups of PRCs, green circles indicate groups of RN neurons, red circles indicate groups of iONR neurons, and blue circles indicate groups of eONR neurons. The rightmost sketch illustrates how the action of RN cells onto eONR cells results into an effective inhibition of the latter.

**Table 1:**
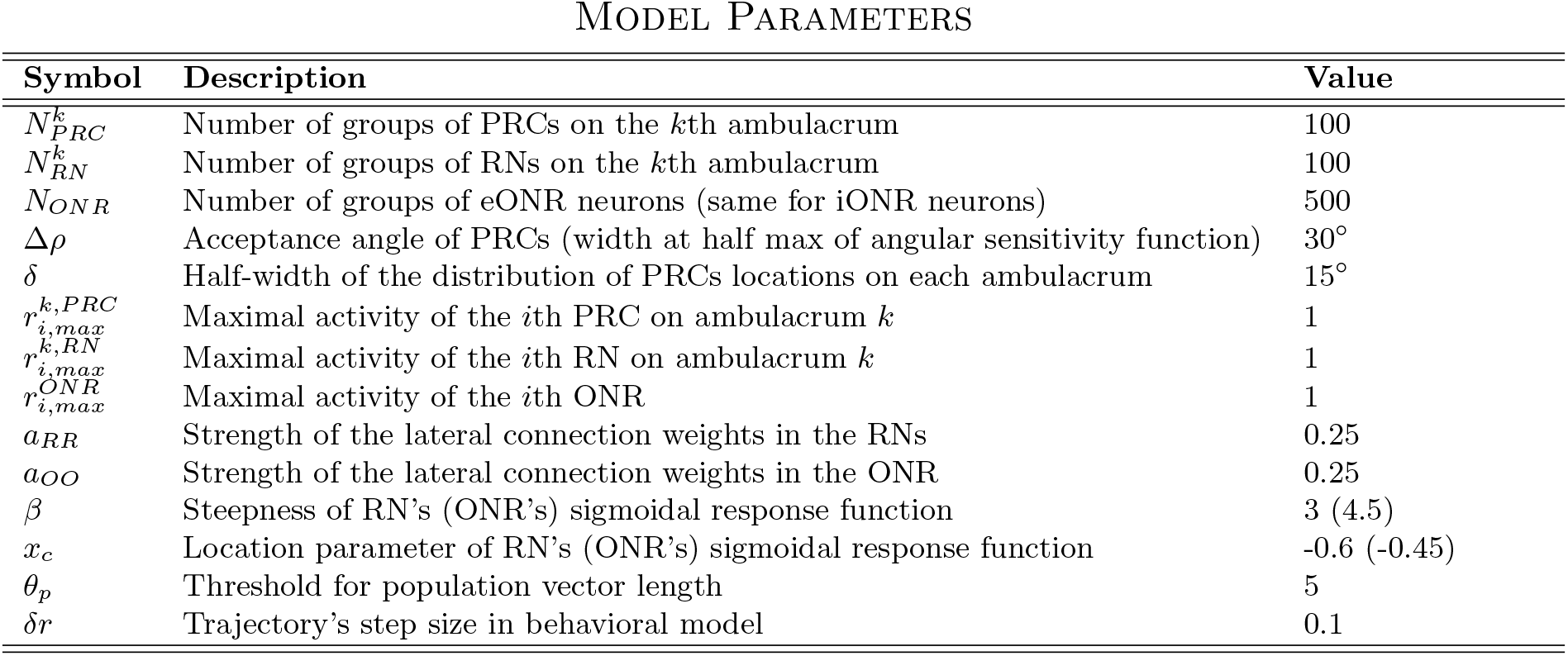
Description and numerical values of the model parameters. See the text for details.

Once activated by light, PRCs inhibit connected groups of RN neurons on the same ambulacrum. RN neurons were assumed to be excitatory, with different RN neuron groups exciting one another. The same groups of RN neurons send excitatory projections to input stage inhibitory cells in the ONR (iONR cells). The latter inhibit the activity of downstream groups of excitatory ONR neurons (eONR cells), that could be interpreted as the output sublayer of ONR. In turn, these groups of eONR neurons excite one another via recurrent connections (Fig. 1B). This allows to integrate information across the five ambulacra, thus mediating a form of decentralized vision.

#### Photoreceptor cells

PRCs are located inside the tube feet protruding along each ambulacrum (Burke et al., 2006; Ullrich-Lüter et al., 2011). The available anatomical description of PRCs, coming mostly from studies in the species *Strongylocentrotus purpuratus* (Agca et al., 2011; Lesser et al., 2011; Ullrich-Lüter et al., 2011), is still incomplete. We consider in the model only the PRCs located at the base of the tube feet, near the test, whose position can be considered fixed in the coordinate system of the sea urchin (see the Supplementary Information, Sec. A.2, for details on the coordinate systems). We did not consider the PRCs located at the tips of the tube feet because, given their lack of screening pigment and their high motility (Ullrich-Lüter et al., 2011), those PRCs cannot contribute to spatial vision (see also the Discussion, Sec. 3.2.1). In the absence of more detailed information about the spatial distribution of PRCs, we assumed a uniform distribution of PRCs in each ambulacrum; different choices, such as a Gaussian distribution, led to similar results (not shown). Since changes along the latitudinal direction are not relevant in our model, we focus again on the longitudinal direction *ϕ* (i.e., from west to east; see Fig. 1A). Along the *k*th ambulacrum, we assume a uniform distribution between of PRCs between *ϕ*^*k*^ −*δ* and *ϕ*^*k*^ + *δ*, where *ϕ*^*k*^ is the position of the center of the ambulacrum (we used *δ* = 15° in the main simulations; see Table 1). Each PRC had an acceptance angle (i.e., the full width at half maximum of the normalized angular sensitivity curve) of Δ*ρ* = 30°. Other values of *δ* and Δ*ρ* could be used and we show later how our results depend on variations of these values (Fig. 6). Our behavioral experiments (Kirwan et al., 2018) suggest acceptance angles in the range 38°-89°. As this estimate takes also into account the distribution width of PRC locations on each ambulacrum, here quantified by *δ*, we chose *δ* = 15° and Δ*ρ* = 30° so as to obtain an ‘effective’ acceptance angle of Δ*ρ*_*eff*_ = Δ*ρ* + 2*δ* = 60° (see also the Discussion, Sec. 3.2.1).

Each PRC’s angular sensitivity curve was modeled as a Gaussian function normalized to have a unitary peak and cut-off at its tails (Land and Nilsson, 2012). We approximated such a function with a cosine function that optimally matches the Gaussian function away from its tails, while naturally vanishing in the corresponding tail regions (Georgopoulos et al., 1982; Salinas and Abbott, 1994) (see Sec. 5.1.2 of Methods for details). The angular sensitivity curves are plotted in Fig. 2A for PRCs in all five ambulacra (each ambulacrum in a different color).

**Figure 2:**
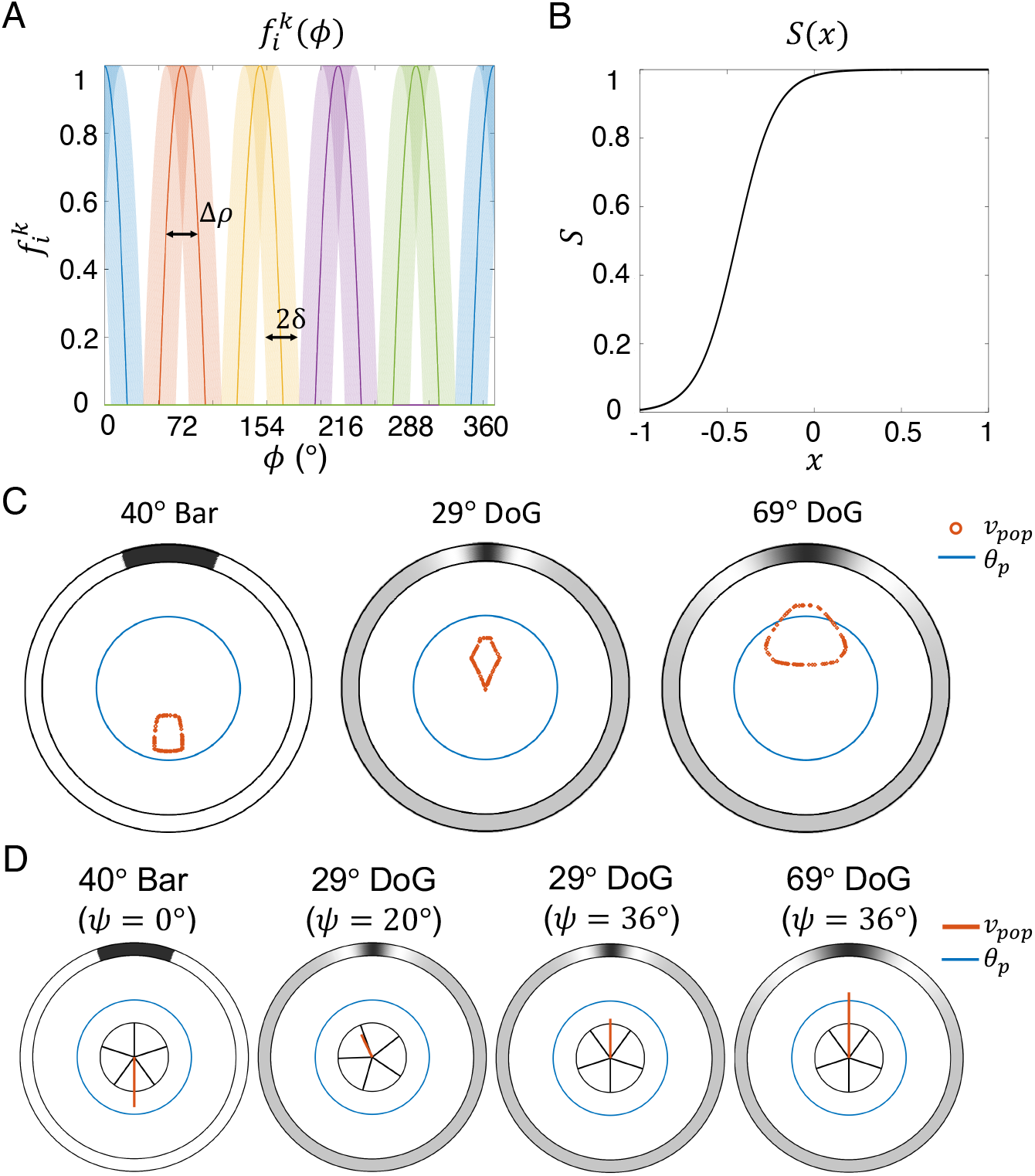
Angular sensitivity curves and population vectors. **A**. Angular sensitivity curves of PRC 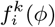 (Eq. 3 of Methods with Δ*ρ* = 30° and *δ* = 15°). Each color represents an ambulacrum. The light shades represent the angular sensitivity curves of all PRCs (uniformly distributed with half-width *δ* in each ambulacrum). The darker lines are examples of single angular sensitivity curves (one example for each ambulacrum). **B**. Plot of sigmoidal function S (Eq. 2) used to model the output of RN and ONR neurons (here shown for the ONR neurons, *β* = 4.5 and *x*_*c*_ = −0.45; see Table 1). Note that activation of PRCs results in a reduced overall input *x*, reducing the output of RN cells (see the text). **C**. Population vectors (readout of eONR cells, see Eq. 11 of Methods) for three stimuli: 40° bar, 29° DoG and 69° DoG (from left to right). Each data point (red circle) corresponds to the tip of one population vector. Different population vectors were obtained by varying the orientation of the sea urchin with respect to the center of the stimulus (here, located at the top of the arena). All orientations (from 0° to 359°) are represented. The large blue circle is the threshold *θ*_*p*_. If for a given orientation the population vector’s length exceeds *θ*_*p*_, visual detection occurs and coherent motion is predicted along the direction of the population vector. **D**. Examples of single population vectors among those in panel C for specific orientations (*ψ*) of the animal with respect to the center of the stimulus (see Suppl. Fig. S1A).

The angular sensitivity curve characterizes the response of a PRC to light coming from a punctiform source located at a given longitudinal angle *ϕ*. The response of the same PRC to a an extended stimulus comprising light coming from all directions was obtained by integrating the stimulus intensity along the angular dimension *ϕ*, weighted by the angular sensitivity curve (see Sec. 5.1.2 of Methods for details). For an animated illustration of PRCs activity in response to stimuli, see the Supplementary Information, Sec. A.4.

#### Radial nerve (RN) neurons

Radial nerve neurons are also distributed along ambulacra (Cobb, 1970; Burke et al., 2006; Formery et al., 2021). In our model, we assumed that they receive inhibitory input from the PRCs (see below). PRCs on ambulacrum *k* project to their target group of RN cells on the same ambulacrum via a (negative) connection weight 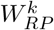. RN cells are also connected to other RN cells on the same ambulacrum *k* via a (positive) connection weight 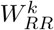 (see Fig. 1B). As illustrated in the figure, we assume that these ‘lateral’ connections among RN neurons are mostly local, i.e., they exist among adjacent groups of RN neurons, although this hypothesis is more a matter of convenience than a crucial ingredient of the model. We also note that these connections may or may not be established via chemical synapses – an issue that has not been settled, not even at the neuromuscular junction (Kawaguti, 1964; Florey and Cahill, 1980), but one that is not essential at our level of description.

The activity of RN neurons in response to input coming from both PRCs and other RN neurons was modeled as a sigmoidal function of the input,

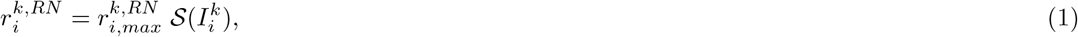

where 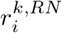 is the output firing rate of the *i*th RN cell on ambulacrum *k*, 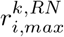 is the maximal firing rate (here, the same for all *i*), and S(*x*) is the sigmoidal function

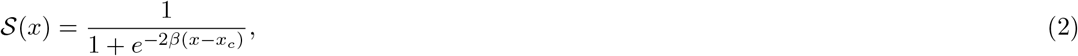

see Table 1 (plotted in Fig. 2B). The input 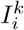 comprises multi-unit activity from PRCs and RN cells targeting RN cell *i* on ambulacrum *k* (these are visualized, in Fig. 1B, by the encircled groups of PRCs and RN neurons targeting the same group of RN neurons). The exact analytical form of the input is reported in the Methods (Eq. 7 of Sec. 5.1.3). Note that since the PRCs provide an inhibitory input to RNs, 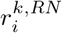 is a monotonically *decreasing* function of the activity of the PRCs: the larger the PRC activity, the smaller the firing rate of RN neurons. In the absence of visual input, the RN neurons maintain a spontaneous firing rate 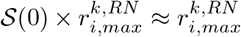 even in the absence of lateral input from other RNs, since the sigmoidal function is saturated at *x* = 0 (see Fig. 2B). This assumption was made to explain the lesion studies reported by Yoshida in the sea urchin *Temnopleurus toreumaticus* (Yoshida, 1966) (see Discussion, Sec. 3.2.2). For an animated illustration of RN neurons’ activity in response to stimuli, see the Supplementary Information, Sec. A.4.

#### Oral nerve ring (ONR)

We assumed that the ONR contains inhibitory (iONR) and excitatory (eONR) neurons, with iONR neurons being the input stage of ONR (Fig. 1B). The RN cells project to iONR neurons via connection weights *W*_*iOR*_, where the connection exists only between specific groups of iONR neurons and RN neurons in their vicinity, i.e., located on the ambulacrum closest to their target group. Groups of iONR neurons, in turn, inhibit groups of eONR neurons in their vicinity via negative connection weights. The overall effect of this connectivity pathway is inhibition of eONR neurons by RN input, and can be described as being mediated by effective (negative) connection weights *W*_*OR*_ (Fig. 1B). In turn, neighboring groups of eONR neurons are connected through (excitatory) weights *W*_*OO*_. These lateral connections allow the ONR to integrate information coming from RNs on different ambulacra. The output of eONR neurons, the firing rate 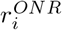, is a sigmoidal function of their input, analogously to the output of RN neurons (see Methods, Eq. 10 of Sec. 5.1.4). The firing rate 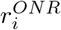 increases due to lateral ONR input and decreases due to RN input. For an illustration of ONR neurons’ activity in response to stimuli, see the Supplementary Information, Sec. A.4.

#### ONR readout and spatial vision

Sea urchins can move towards or away from specific visual stimuli that they are able to detect (Yoshida, 1966; Blevins and Johnsen, 2004; Yerramilli and Johnsen, 2010; Kirwan et al., 2018; Al-Wahaibi and Claereboudt, 2017). In our model, the direction of movement was determined by pulling together the activity of eONR neurons to produce a specific vectorial readout named ‘population vector’. The population vector is built by assigning a ‘vote’ to each eONR neuron according to its own ‘preferred direction’, which was defined as the direction of a narrow stimulus causing the maximal increase in activity in the cell, and is highly correlated with the cell’s angular location (see Methods for details, Sec. 5.1.5). When an eONR is active, its firing rate ‘votes’ for its preferred direction. The directions associated to different eONR neurons are then added up as vectors, and the resulting vector (the population vector) is compared to the threshold *θ*_*p*_. If the length of the population vector exceeds the threshold, the urchin detects a visual stimulus as coming from the same direction as the population vector. The longer the population vector, the more reliable the detection.

The population vectors in the presence of the three spatially extended stimuli utilized in the experiment of (Kirwan et al., 2018) are shown in Fig. 2C for all possible orientations of the animal with respect to the center of the stimulus (see Fig. 3 for a description of the stimuli). This figures shows that a stimulus may or may not be visible depending on the relative positions of the sea urchin and the stimulus. Therefore, visible stimuli are those for which there is a subset of orientations for which the population vector exceeds *θ*_*p*_ (blue circle in figure). For animations of this figure as the animal and stimulus change their relative orientation, see the Supplementary Information, Sec. A.4.

**Figure 3:**
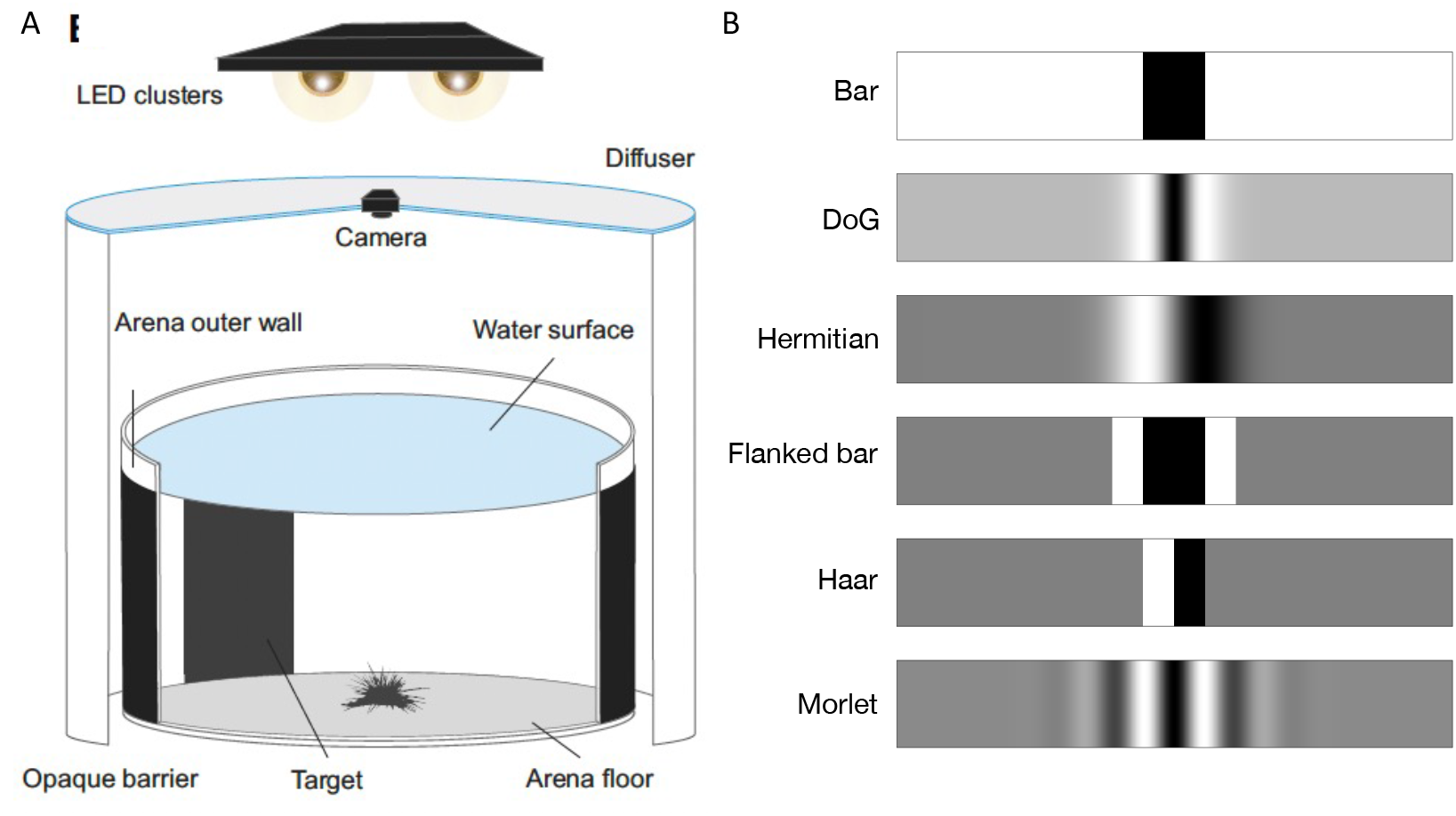
Experimental setting and stimuli. **A**. Behavioral experiment setting. The stimuli were attached on the outer wall of the arena. At the beginning of each trial, the animal was positioned at the center of the arena with a random orientation. **B**. Examples of the six stimuli used in this work with *ϕ*_*stim*_ = 40° (mathematical definitions in Suppl. Table S1). The ink value of black and white are 1 and 0, respectively. Panel A adapted from (Kirwan et al., 2018).

### 2.2 Model prediction of object taxis in *Diadema africanum*

The model developed in the previous subsections was tested on the visually-guided behavior of *D. africanum* in taxis experiments performed by Kirwan et al. (2018). In the experiments, individuals of this species were located at the center of an arena with printed stimuli attached to the outer wall of the arena in a way to completely surround it (a schematic illustration of the arena is shown in Fig. 3A). Three different stimuli were used in the experiments: a 40° bar stimulus and two stimuli obtained from taking the difference of two Gaussian functions (‘difference of Gaussians’, or DoG), each with a darker center subtending a 29° and a 69°, respectively. These stimuli are shown as the top two stimuli in Fig. 3B and were used to match the model to the experimental data. The remaining stimuli shown in Fig. 3B were used to make novel model predictions on future experiments (see the Supplementary Information, Sec. A.2, for the mathematical characterization of all the stimuli).

The behavioral results found by (Kirwan et al., 2018) are shown in Fig. 4A. In summary, the sea urchins moved randomly in the presence of a homogeneous stimulus of constant intensity that was used as control (Fig. 4A, top left panel). Similarly results were obtained in the presence of a 29° DoG stimulus (Fig. 4A, top right). Some coherent movement towards a 40° bar stimulus was detected, however its statistical significance was unclear (Fig. 4A, bottom left); and finally, there was clear (significant) movement towards to center of a 69° DoG stimulus (Fig. 4A, bottom right).

**Figure 4:**
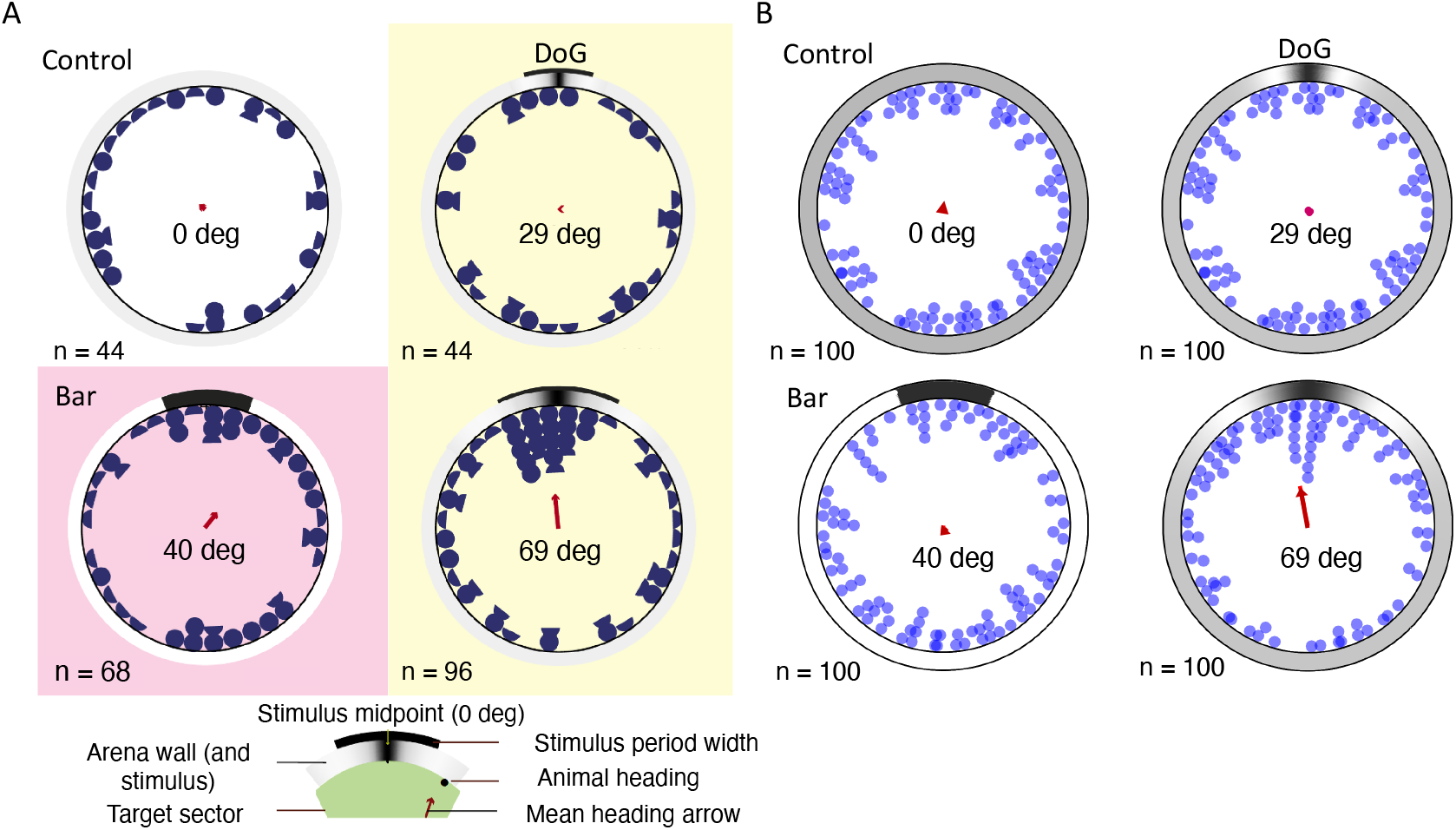
Behavior of model and *Diadema africanum* in the presence of 4 different stimuli. **A**. Experimental results from (Kirwan et al., 2018). Each blue semicircle represents the final position of the animal at the end of one trial, while a full blue circle represents the final position of two animals. Although all animals reached the wall of the arena, final identical positions were stacked to show all the data. The stimuli used were a stimulus with uniform intensity in all directions (control; top left panel); a 29° DoG stimulus (top right); a 40° bar (bottom left); and a 69° DoG (bottom right; see Fig. 3 and Table S1 for details about the stimuli). In each panel, the stimulus is shown on the outer wall of the arena (large circle). The red arrow at the center of the arena is a measure of aggregate directional movement across each cohort of subjects (see below). **B**. Model results in simulations of the same tasks shown in panel A. Each blue circle corresponds to one predicted final position across a cohort of 100 animals with random initial orientations with respect to the center of the stimulus. Here, the final position was inferred from the population vector induced by the stimulus when the animal was at the center of the arena (see Fig. 2C-D). In both panels, the red arrow at the center of each plot is circular mean vector of the final positions (see Methods, Eq. 14). The only significant circular mean vector was obtained with the 69° DoG stimulus: *P* = 0.013 (V-test) and *P* = 0.042 (Rayleigh test), in agreement with the analogous results by (Kirwan et al., 2018) (*P* > 0.5 for the remaining stimuli). Main parameters: Δ*ρ* = 30*°, δ* = 15*°, θ*_*p*_ = 5 (see Table 1 for all other parameter values). Panel A adapted from (Kirwan et al., 2018).

Significant aggregate behavior towards the 69° DoG stimulus was not merely due to a larger number of subjects used in this task (*n* = 96) compared to the same task with the other two stimuli (see (Kirwan et al., 2018) for details). The red vector at the center of the arena is the circular mean vector (see Methods Eq. 14). The direction of this vector indicates the circular mean orientation of all bearings, while the length of the arrow indicates the ‘mean resultant length’, a measure of the concentration of the points along the direction of the arrow (the longer the arrow, the more coherent directional motion in the direction of the arrow compared to the uniform distribution; a short arrow indicates random direction of movement (Kirwan et al., 2018)).

The experimental results are accurately predicted by the model, as shown in Fig. 4B. Model sea urchins with random orientations were located at the center of the arena, and the final position in each case was inferred from the initial value of the population vector (the collective readout of eONR activity). For population vectors of length below a threshold *θ*_*p*_ = 5, a random final position was inferred, otherwise the final position was drawn from a Gaussian distribution with the same mean as the population vector. The distribution of final positions in Fig. 4B is due to random initial orientations across different bearings (i.e., visual detection depends on the relative position of animal and stimulus) and reflects the distribution of population vectors shown in Fig. 2C (or a circularly uniform distribution if the populations vectors are all smaller than the threshold; see Methods, Sec. 5.1.5, for details). This ability for spatial vision is the result of integrating light information coming from all PRCs distributed on the animal’s body, mediated by neural activity in each RN, and finally integrated in the ONR.

As shown in Fig. 2C, the population vector is always below threshold for the 29° DoG and the 40° bar, predicting random motion as confirmed in Fig. 4B. For the 69° DoG stimulus, the population vector length exceeds the threshold when the region of the animal’s body between two ambulacra is facing the ‘target region’ of the stimulus (roughly, the dark region flanked by the white maxima; see Fig. 2D, rightmost plot). This means that, averaging across all initial orientations, there will be detectable motion towards the target, as confirmed in Fig. 4B (bottom right). The simultaneous activation of the activities in PRCs, RNs and ONR neurons, together with the resulting population vector readout in the ONR, can be appreciated in the animations presented in the Supplementary Information (Sec. A.4).

### 2.3 Dynamic model of object taxis

In the previous section we have inferred the final position of the animal based on its ability to detect the stimulus when located at the center of the arena. This method assumes that the initial population vector not only establishes whether the stimulus has been detected, but it is also a proxy of the final position of each bearing. As the animal moves towards the target, however, stimulus detection will change due to a change in the relative position of stimulus and animal. Such relative position will change at every movement and can in turn produce adjustments in movement direction. It is therefore of interest to have a model of visually-driven, step-by-step movement. This would also allow a better comparison with the the experimental results shown in Fig. 4A, where the final position was determined based on the projection, on the arena wall, of the line connecting the center of the arena with the location of the animal at one quarter of the radius of the arena (Kirwan et al., 2018).

To include a basic mechanism of movement, we discretized time into small steps and predicted the movement direction of the next step based on i) the stimulus as detected by the animal at the current position, and ii) the direction of previous movement (see Sec. 5.1.6 of Methods for details). Fig. 5 shows the simulated trajectories for the same four stimuli shown in Fig. 4B (see the Supplementary Information, Sec. A.5, for full animations). Each colored curve is the trajectory in one out of 100 trials and the final position of the center of the body is marked by dot of the same color (radially projected as a grey dot on the arena wall). The final position of the body center was set at 1*/*4 of the distance between the wall and the center of the arena to take into account the long spikes of *D. africanum*, so that the edge of the body reaches the wall when the center is at such distance from it.

**Figure 5:**
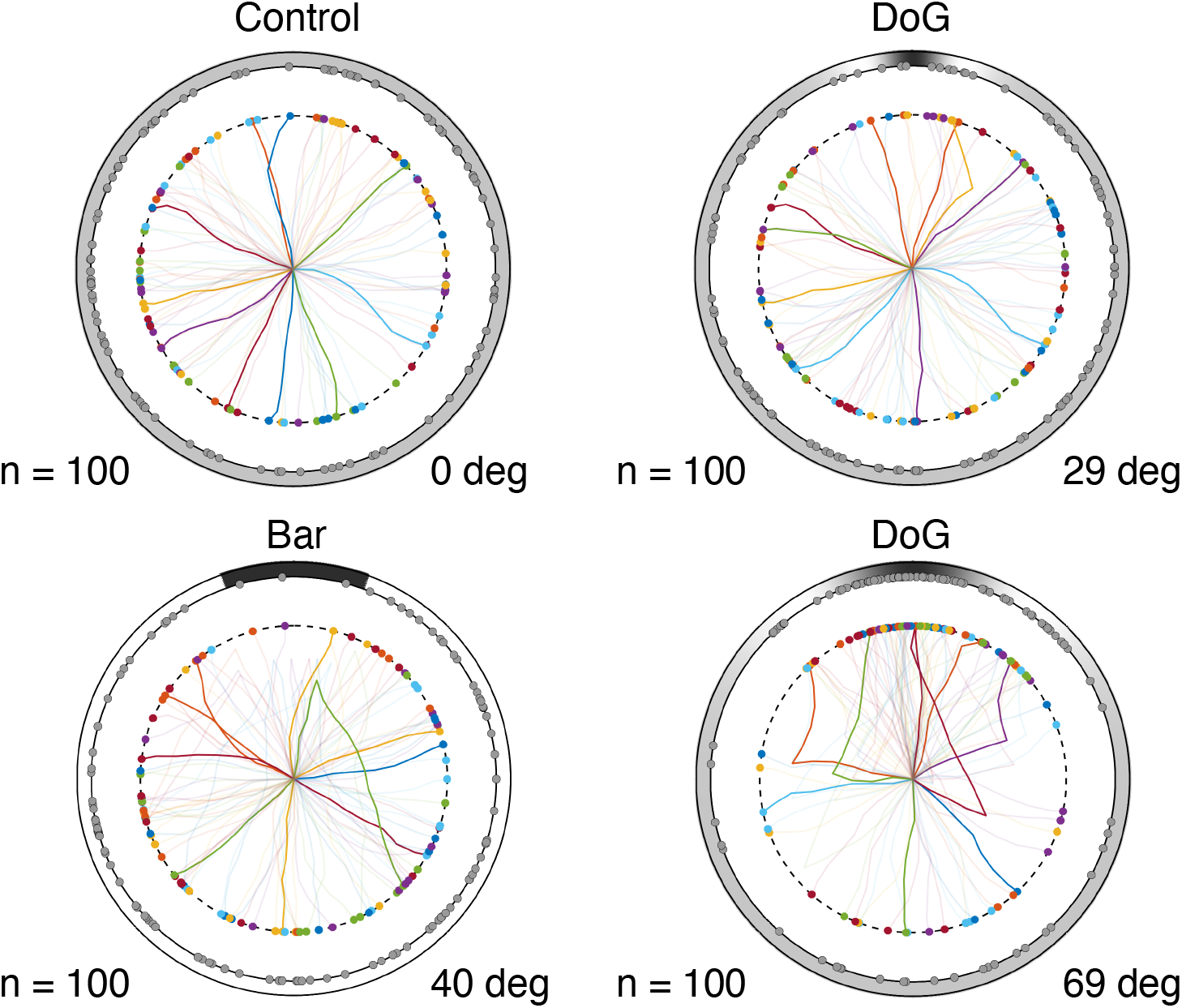
Simulated trajectories of 100 bearings under control, 40°bar, 29° DoG and 69° DoG stimuli. The outer circle represents the wall of the arena with the stimulus on it. Animals were placed at random orientations at the center of the arena and moved according to the behavioral model described in section “Model of visually-induced movement”. The colored lines represent the trajectories covered by the center of each animal starting from the center of the arena and ending at the color-matched dot. The grey dots on the arena’s wall are the radial projections of the colored dots and represent the points where the animal’s body hits the wall. See Table 1 for model parameters.

The distributions of predicted final positions agree well with those in Fig. 4B, suggesting that the initial movement produced at the center of the arena is, on average, a valid proxy for the final position reached using this simple model of behavior. Under control, the trajectories started off a random direction and did not change direction except for small random changes around the previous direction. For the 29° DoG, initial movement was random but it would occasionally turn towards the stimulus. This happened when the location of the animal was such that light coming from the white maxima of the DoG struck one ambulacrum, causing a strong activation of PRCs at that location. For the 40° bar, the animal moved randomly at first but changed movement direction later in several trials. These changes occurred close to the stimulus target (the black region), which became wider in reference to the size of the animal as the latter approached the arena. On the opposite side, the bright region of the stimulus became narrower so that fewer ambulacra were activated. This resulted in a longer population vector and a change of movement direction towards the opposite side (this is due to a reduced cancellation of visual information that would occur when uniform light comes from most directions). Compared to the ‘static’ model of Fig. 4B, we noticed a smaller fraction of animals reaching the wall near the target region, although this caused no appreciable difference in the circular mean vector. Finally, for the 69° DoG, the animal would turn towards the the target region when the latter faced directly one ambulacrum. In this case, due to the 69° arc subtended by the stimulus, each of the two white maxima of the stimulus would face one ambulacrum, producing a large population vector in the direction of the target region (see the rightmost plot in Fig. 2D). As a result, there was a significant concentration of final positions near the target stimulus, as also observed in Fig. 4B (bottom right).

### 2.4 Effect of location and acceptance angle of PRCs on spatial vision

The results shown so far were obtained assuming a longitudinal distribution of PRCs with half-width *δ* = 15° and acceptance angle (the half-width of the angular sensitivity function) of Δ*ρ* = 30°. Together, this results in an effective acceptance angle of 60° (Fig. 2A). Varying either *δ* or Δ*ρ* can affect detection acuity (the minimum angular width of a visual stimulus which can be detected). For example, the reason why the model fails to detect the 29° DoG stimulus is the small arc length separation of the white maxima of the stimulus (the latter can only activate PRCs located on one ambulacrum; see Fig. 2CD).A larger distribution width or a larger acceptance angle, however, could allow the activation of PRCs on two ambulacra, as it occurs in the 69° DoG. Is our model robust to variability of *δ* and Δ*ρ*? In other words, what range of values can be allowed for these parameters so that the model still captures the results of Fig. 4A?

To answer this question, we computed the maximal length *v*_*max*_ of the population vector, Eq. 11, across all possible orientations. This was done for all values of Δ*ρ* between 15° and 90° and *δ* between 5° and 20°(Fig. 6). If, for a given pair (Δ*ρ, δ*), one has *v*_*max*_ > *θ*_*p*_, then the stimulus can be detected (a more realistic criterion is to require a finite range of initial orientations for which the population vector length exceeds *θ*_*p*_). Note that the range of Δ*ρ* values used here contains our previously estimated range based on contrast thresholds between 5 and 20% (Kirwan et al., 2018).

**Figure 6:**
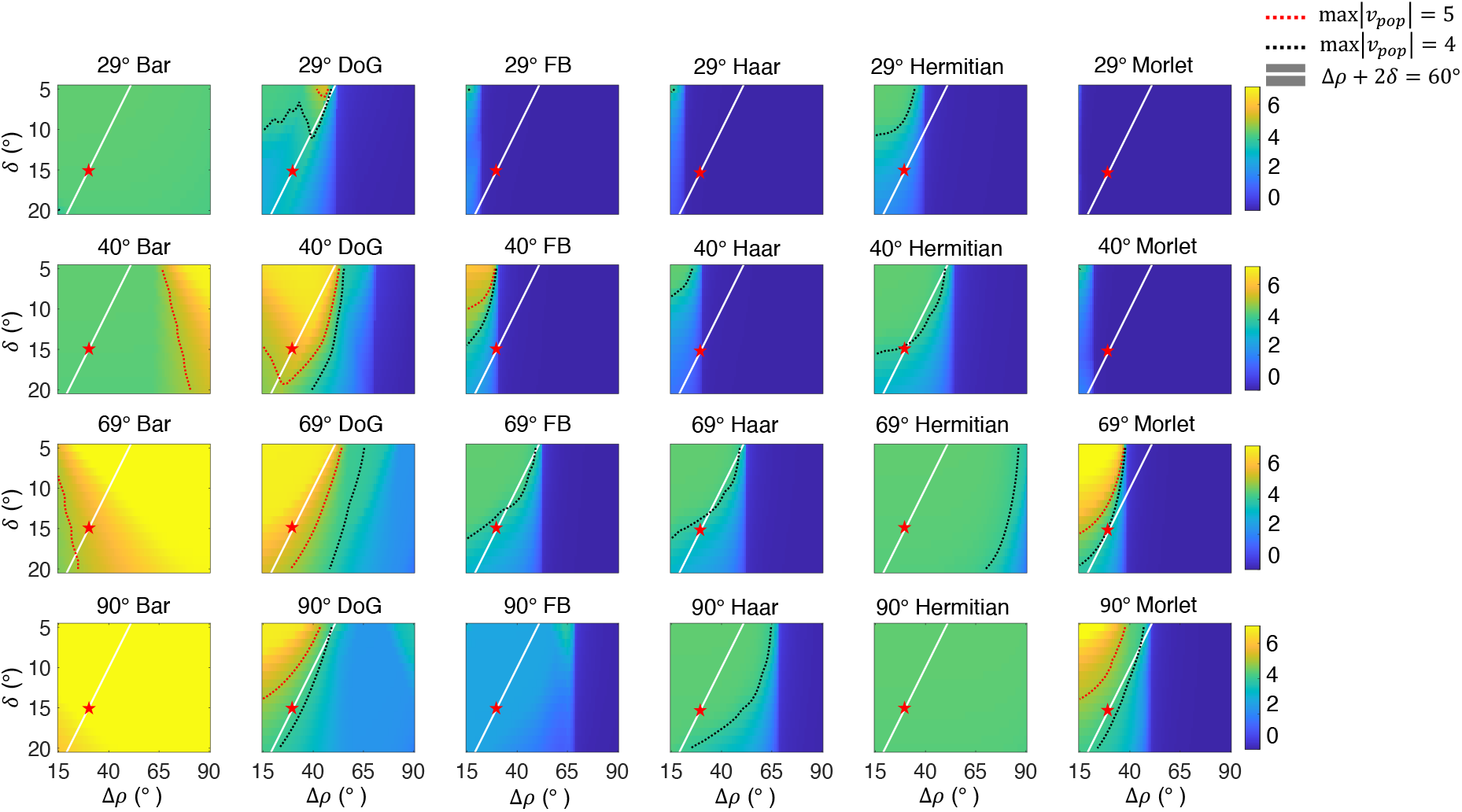
Effect of acceptance angle and location of PRCs on the model’s spatial vision. Each panel shows a heat map of *v*_*max*_, the maximal length of the population vector across initial orientations of the animal, for a given stimulus and a given pair of values for Δ*ρ* and *δ*. Each column shows the same stimulus for different arc widths of the target region (i.e., for different *ϕ*_*stim*_, see Table S1), while each row shows the same *ϕ*_*stim*_ across different stimuli. If *v*_*max*_ < *θ*_*p*_, the animal cannot detect the stimulus from any orientation. The red dotted line is the contour line where *v*_*max*_ = *θ*_*p*_, while the black dotted line is the contour line where *v*_*max*_ = 4. The white line is the collection of points with Δ*ρ* + 2*δ* = 60°. In each plot, Δ*ρ* ranged between 15° and 90° and *δ* ranged between 5° and 20°. Red stars mark the point (Δ*ρ, δ*) = (30°, 15°), the parameter values used in the main simulations. *DoG*: Difference of Gaussians; *FB*: Flanked bar.

Fig. 6 shows the value of *v*_*max*_ for different values of Δ*ρ* and *δ* for a wide range of stimuli, beyond those used in our previous experimental study (Kirwan et al., 2018). In each plot, the red dotted line is the contour line where *v*_*max*_ reaches the boundary defined by the threshold (*θ*_*p*_). The animal is able to detect the stimulus for parameter values (Δ*ρ, δ*) inside the yellow region delimited by the red dotted line.

The red stars mark the pair of parameter values (Δ*ρ, δ*) = (30°, 15°) used in Fig. 4B. The point defined by this pair is located outside the yellow region for the 40° bar and 29° DoG, whereas it is inside the yellow region for the 69° DoG stimulus. This implies an animal with (Δ*ρ, δ*) = (30°, 15°) can detect the 69° DoG, but not the 40° bar and 29° DoG, as already established. Fig. 6 also shows that our results are robust to some degree of parameter variation, in the sense that they hold in a finite region of parameters space around the point (Δ*ρ, δ*) = (30°, 15°).

Our model makes other interesting predictions. For example, it predicts that the impact of Δ*ρ* on spatial vision depends on the stimulus. For some stimuli (e.g., the bar) detection improves for larger Δ*ρ*, while for other stimuli (e.g., the DoG and the Morlet wavelet) detection improves for smaller Δ*ρ*. Furthermore, the model can be used as a guide for choosing probing stimuli in future behavioral experiments. For example, the model predicts that a 69° bar stimulus should be easily visible, whereas a 69° Flanked bar, Haar wavelet, or 1st Hermitian wavelet stimulus should be hard to detect from any initial orientation. We emphasize that these predictions – which can be tested in experiment – depend on the neural model built in this study, and not just the properties of the PRCs.

Finally, note that in Fig. 6 we have assumed an equal acceptance angle Δ*ρ* for all PRCs, which is unreasonable. To address this issue, we performed the same analysis using a random distribution of Δ*ρ* values across PRCs in each ambulacrum. As shown in the Supplementary Information (Sec. A.6), a random distribution of Δ*ρ* values does not alter the general picture shown in Fig. 6 (compare Fig. 6 with Fig. S4).

## 3 Discussion

### 3.1 A model of decentralized vision in sea urchins

In this work we propose a model of decentralized vision in sea urchins based on general anatomical features of their nervous system and assuming that light information is captured by photoreceptor cells (PRCs) distributed over the animals’ dermis around the ambulacra. PRCs have been found on the tube feet of sea urchins (Agca et al., 2011; Lesser et al., 2011; Ullrich-Lüter et al., 2011), which emerge from the test of the animal along the five ambulacra. The model considers the nervous system of the animal in some degree of detail, and provides a mechanistic model involving all stages of the system, from PRCs to neural activity in the ONR. As information flows from PRCs to RN neurons to ONR neurons, the model can be abstracted as comprising three major layers. The model has feedforward connections between the layers (PRCs to RN neurons and RN neurons to ONR neurons) as well as recurrent connections among neurons in the same RN, and within neurons of the ONR (Fig. 1B). PRCs on the tube feet of the animal, once activated by light, inhibit RN neurons in the same ambulacrum. In turn, RN neurons inhibit excitatory neurons in the ONR (eONR). Lateral recurrent connections in the ONR allow to integrate the information coming from all ambulacra. The activity of ONR is read out as a ‘population vector’. The population vector collates the preferred directions and the activity of all eONR neurons in the presence of a specific visual stimulus. When the length of the population vector is large enough to exceed a threshold, enough evidence has been accrued about the location of a visual stimulus, and movement towards said location occurs with probability proportional to the length of the population vector. We discuss below the feasibility of all these putative mechanisms in light of the available experimental evidence.

The model explains in quantitative detail the behavior of *D. africanum* found in the taxis discrimination task of (Kirwan et al., 2018), where taxis was found clearly only in 69° isoluminant DoG stimuli, but neither in narrower DoG nor in a 40° bar stimulus. The latter finding was somewhat surprising given that there is a much greater local contrast in the case of the bar stimulus and greater amplitude at low spatial frequencies (see Fig. 4 of (Kirwan et al., 2018)). Our model provides a mechanistic explanation of these results based on generic anatomical features of the nervous system of sea urchins combined with a specific mechanism of visual integration and readout at the level of ONR neurons.

### 3.2 Assumptions of the model

In building our model, we have considered the available experimental data on sea urchins anatomy and behavior, and turned to other systems when the experimental knowledge was insufficient. This has led us to combine elements from both invertebrate and vertebrate nervous systems, some of which inspired by mammalian physiology. Examples of invertebrate structures are the ganglia-like clusters of RNs connected to clusters of PRCs, and nearby clusters of RN nerves connected ‘laterally’ (see Fig. 1B) (Hyman, 1955; Matheson, 2002). This local arrangement is compatible with the fact that axons of echinoderm neurons are normally small and unmyelinated, and are bundled in packages with a parallel arrangement (Ortega and Olivares-Bañuelos, 2020). Examples from vertebrate animals include sigmoidal ‘tuning curves’ to characterize the response of RN and ONR neurons (Eq. 2). Neurons with similar tuning curve properties are found e.g. in cats and primates and include primary and secondary visual neurons (Albright, 1984; Maunsell and Van Essen, 1983), middle temporal neurons (Albright, 1984; Britten et al., 1993), parietal neurons (Fanini and Assad, 2009), motor cortical neurons (Georgopoulos et al., 1982), and so on, suggesting perhaps that tuning curves, as a way to relay sensory information, may be widespread across different nervous systems, echinoderms included. Similarly, the idea of reading out the activity of ONR neurons via population vectors was borrowed from landmark studies in motor and premotor cortex of primates (Kalaska et al., 1983; Georgopoulos et al., 1982).

While the existence of analogous properties in sea urchin neurons awaits experimental demonstration, we included these structures in our model for a number of reasons. First, some evidence in the literature suggests that we may be on the right track. For example, studies in other echinoderms such a brittle stars (Clark et al., 2019) indicate that bidirectional connections between RNs (similar to our model’s lateral connections) are necessary for coordinated locomotion. However, the analogy with better studied marine invertebrates is often of very limited help. For example, starfish orienting in an odor plume lead with the rays facing the odor source and change leading ray multiple times during the approach to the source (Dale, 1999), whereas we did not reliably observe similar changes in our taxis experiments (Kirwan et al., 2018). Finally, it seems that some neural elements are indeed common to the mammalian and echinoid nervous systems. For example, (Pentreath and Cobb, 1972) have reported cholinergic and dopaminergic neurons in the central nervous system of echinoderms, with acetylcholine bound to synaptic vesicles morphologically similar to those present in the mammalian brain.

To summarize, our modeling choices were informed by arguments of plausibility and generality whenever the experimental information was inadequate. Our most salient choices are discussed in more detail below.

#### 3.2.1 Model of tube feet photoreceptor cells

PRCs at the base of tube feet are located along the ambulacra, but their detailed distribution is not known. We have therefore chosen a uniform distribution of locations that spans 30°longitudinally. This feature of the model does not present a limitation, for two reasons: i) other distributions of PRC locations (such as a Gaussian distribution) result in a quantitatively equivalent model (not shown); ii) the model’s behavior is robust to variations of parameter values such as location spread, *δ*, and acceptance angles, Δ*ρ*, of single PRCs (Fig. 6).

The values chosen for these parameters reflect the position of the PRCs on the tube feet and other factors, such as depressions in the test that can hold space for r-opsin expressing PRCs while also screening light reaching the PRCs (Ullrich-Lüter et al., 2011), or the shading activity of opaque spines (Woodley, 1982; Blevins and Johnsen, 2004; Yerramilli and Johnsen, 2010). Our behavioral experiments (Kirwan et al., 2018) suggest acceptance angles in the range 38°-89°. As this estimate takes also into account the distribution width of PRC locations on each ambulacrum, we chose *δ* = 15° and Δ*ρ* = 30° so as to obtain an ‘effective’ acceptance angle of Δ*ρ*_*eff*_ = Δ*ρ* + 2*δ* = 60°.

We note that although a sizable density of PRCs is found on the tips of the tube feet, the latter lack any associated screening pigment and are highly motile (Ullrich-Lüter et al., 2011). As a consequence, the PRCs located on the tube feet disks display continuously changing spatial properties and cannot provide the basis for spatial vision, unlike the PRCs at the base of the tube feet. Moreover, in a recent analysis of scRNA-seq data of early developmental stages of the sea urchin *Paracentrotus lividus*, different gene expression profiles were found for two sub-clusters of r-opsin expressing PRCs (Paganos et al., 2022). Those findings might reflect different functions of the two PRC clusters located, respectively, at the tip and base of the tube feet that had been initially reported in (Ullrich-Lüter et al., 2011).

Finally, although sea urchins can presumably sense light gradients along the latitudinal (vertical) direction, we did not consider this possibility in our model, since our stimuli were vertically homogeneous and locomotion occurred always on a horizontal plane. We also grouped cells with similar properties in each layer (PRC, RN or ONR) and averaged their behavior (input and output), so that we could consider a homogeneous group of neurons as the elementary processing unit in our model. This type of coarse-grained approach is typically successful in systems with redundant numbers of elements and it allows to focus on the essential features of the anatomy.

#### 3.2.2 Double inhibition of oval nerve ring neurons

In the model, light-induced excitation of eONR neurons occurs by double inhibition: PRCs inhibit RNs, which in turn project to iONR neurons which inhibit eONR neurons. In this model, the more the RNs are inhibited by light, the larger the response of target ONR neurons leading to stimulus detection. Although this is not the only possible mechanism compatible with our results (double excitation would work just as well), we favored double inhibition over double excitation because the former makes our model compatible with experimental findings reported in other species of sea urchins. Inhibition by light was invoked by Millott and Yoshida (1960) to explain the shadow reaction of *Diadema antillarum Philippi*, and by Yoshida (1966) to explain experiments performed by Yoshida and Kobayashi in the sea urchin *Temnopleurus toreumaticus* (see our summary below). The putative inhibition of RN neurons in response to light is also reminiscent of the ‘off’ response of isolated RNs observed in the sea urchin *Diadema setosum* (Takahashi, 1964), and in general, our model’s assumption that excitatory RN neurons are inhibited by a stimulus resonates with the notion, based on early electrophysiological experiments, that RNs are a locus for interaction between excitation and inhibition (Millott and Okumura, 1968).

In the experiments described in (Yoshida, 1966), four ambulacra were surgically removed and the animal placed in dim light (under photographic safelight). Under these conditions, the majority of animals moved in the direction of the surgically operated area. Assuming positive phototaxis as the most likely behavior in this experiment (wherein the animal moves towards the light source), this finding could be explained if RN neurons inhibited ONR neurons: the removal of RNs would remove an inhibitory factor, increasing the activity of ONR neurons close to the operated area, and producing locomotion in that direction. In a related experiment, a light source was positioned outside of the animal facing the intact RN, and movement towards the light source usually ensued. This is consistent with inhibition of the intact RN by the photo-stimulated PRCs. This inhibition must cause a stronger excitation of the ONR neurons near the intact site, compared to the excitation caused by the absence of RNs on the (operated) opposite site.

When the light source was located *internally* and near the surgically operated site, locomotion in the direction of the intact RN was reduced but was still more likely than locomotion in the opposite direction. This may be because RNs can sense light directly, although to a smaller degree than when stimulated directly by PRCs. In our double inhibition model, this smaller light detection causes a smaller inhibition of the intact RN and therefore a smaller disinhibition of ONR neurons near the intact side, explaining (i) the preferential locomotion towards the intact side, but also (ii) the presence of locomotion towards the opposite side (caused by the absence of RNs on that side). Analogous experiments with two intact RNs (instead of one) led to analogous conclusions (Yoshida, 1966).

By adopting a double inhibition model, we can quantitatively explain the locomotion experiments in *D. africanum* while also capturing, at least qualitatively, Yoshida’s observations in surgically operated sea urchins. In this sense, the model makes strong predictions on the patterns of connections between PRCs, RN neurons and ONR neurons.

In the same article, Yoshida reported another finding that supports the integrative role ascribed by our model to the ONR. Specifically, transecting one RN near the ONR resulted in lack of motion in the direction of the transected nerve, when the animal was placed between two light sources facing each other. Thus, the ONR may play a key role in integrating visual stimuli from RNs. In our model, we have specified a possible mechanism of integration (discussed next) that can explain the results of Kirwan et al. (2018).

#### 3.2.3 Model of visual detection and movement

In the absence of more detailed experimental evidence, we have chosen to follow a principle most precisely formulated in neuroscience (Pouget et al., 2000), according to which sensory and motor functions can benefit from pulling together the activity of many neurons in key areas. In our case, pulling together the collective activity of RNs and ONR neurons determines the strength of visual detection and the stimulus direction. The specific mechanism we adopted makes use of the population vector, which is a natural and general construct for reading out the activity of populations of neurons.

We prefer not to commit to more complex interpretations of the population vector. For example, while the length of the population vector clearly reflects the animal’s ability to detect the stimulus, it may also incorporate the animal’s *motivation* to move towards (or away from) the stimulus, once detected. Therefore, resolving power could be greater than the level exhibited. Also, the same model could be used to move towards the light (as assumed here) or away from it, by simply reversing the meaning of the population vector’s direction. Thus, while the readout of the ONR neurons’ activity could represent additional variables related to motivation or other determinants of the behavior, we feel that it is premature to commit to any of them due to the lack of the necessary experimental data. At this stage, the population vector is best understood as the link between the neural activity produced by the nervous system of our model, and the observed orienting behavior of *D. africanum*.

### 3.3 Questions for future experiments

It is an important question to determine experimentally to what degree our anatomical assumptions hold. Much information is lacking on the nervous system of Echinodermata, partly due to technical difficulties (Ortega and Olivares-Bañuelos, 2020). In the absence of more detailed information, our assumptions allow an explanation of vision in *D. africanum* – at least pertaining to the experiments performed in (Kirwan et al., 2018) – and allow to make clear predictions for future experiments. In turn, the model presented here can be modified and improved on the basis of new experimental evidence, which may alter its functionality to different degrees. To this aim, future anatomical, morphological and behavioral experiments should help clarify issues such as the impact of PRCs input onto clusters of RN neurons; the nature of neural connectivity, including the details of the excitatory vs. inhibitory action exerted by clusters of RN and ONR neurons on their target structures; the electrophysiological response of the same neurons to luminous stimuli (their ‘tuning’ properties); and the detailed nature of ONR integration of RN input. Furthermore, the aggregate response of ONR neurons to luminous stimuli could reveal the details of the mechanism used by these organisms to produce visually-driven behavior (here modeled via the population vector). Finally, comparison of our predictions with behavioral experiments conducted with various stimuli as shown in Fig. 6, could give indirect information on the distribution of PRCs, their (effective) angular sensitivity, and more in general on the architecture of the model upon which the predictions are based.

## 4 Acknowledgments

This work was supported by a Research Grant from the Human Frontiers Science Program (Ref.-No: RGP0002/2019). We thank Drs. Esther Ullrich-Lüter and Carsten Lüter for very useful discussions, Drs. Esther Ullrich-Lüter and Michael Bok for a careful reading of the manuscript, and Dr. Esther Ullrich-Lüter for expert guidance on the literature on echinoderms.

## 5 Methods

### 5.1 Details of the model

#### 5.1.1 Stimuli

In the original experiments of (Kirwan et al., 2018) analyzed here, the printed patterns surrounding the arena were greyscale printed images consisting of dark regions set against a lighter background (shown in Fig. 3B of the main text). The stimuli were uniform in the vertical plane but varied in the horizontal plane along the longitudinal direction *ϕ* in the coordinate system of the arena. This variation was described by a function *X*(*ϕ*) which was the input to the model PRCs (described in detail in the Supplementary Information, Sec. A.2). The main patterns used in the experiment were a 40° bar stimulus and the difference of Gaussian functions (DoG) subtending either 29 or 69 degrees. In the bar stimulus, a region of homogeneously black stimulus subtending 40°was presented against a white background. In the DoG stimulus, the center of the stimulus was maximally dark, but of increasing reflectance towards the periphery of the stimulus (on the horizontal axis) and reaching the maximum achievable reflectance before darkening into the grey background. These stimuli were used in the model to obtain a match between model and experiment. The remaining stimuli shown in Fig. 3B were used to make model predictions for future experiments. All stimuli were isoluminant with respect to the remainder of the patterns due to the lighter regions flanking the stimulus, i.e., it is not possible to detect the stimulus by simply comparing the radiance profile of different parts of the arena from the centre without having a spatial resolution equivalent to the arc subtended by the stimulus itself. Full details about the stimuli can be found in Sec. A.2 of the Supplementary Information.

#### 5.1.2 Photoreceptor cells

Each PRC’s angular sensitivity curve was modeled as a Gaussian function normalized to have a unitary peak and cut-off at its tails (Land and Nilsson, 2012). We approximated such a function with a cosine function that optimally matches the Gaussian function away from its tails, while naturally vanishing in the corresponding tail regions (Georgopoulos et al., 1982; Salinas and Abbott, 1994):

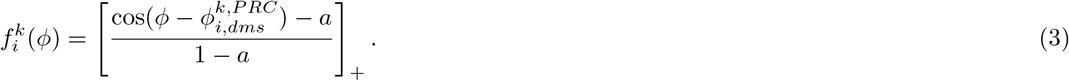

In this function, 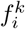 is the response of the *i*th PRC on ambulacrum *k* from punctiform light coming from a longitudinal angle *ϕ*; *a* = 2 cos(Δ*ρ/*2) − 1, where Δ*ρ* is the acceptance angle of the PRC; the symbol [·]_+_ means rectification ([*x*]_+_ ≥0 for *x* ≥0, [*x*]_+_ = 0 otherwise); and 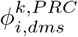 is the direction of maximum sensitivity of the *i*th PRC on ambulacrum *k*. Note that in Eq. 3 only the difference of two angles matters, which is independent of the coordinate system chosen. In the following we consider the angles given in the coordinate system of the sea urchin, which is simply the coordinate system of the arena rotated by a fixed angle *ψ* (the angular distance of the first ambulacrum from the center of the stimulus). For simplicity, we assume that light is screened so that 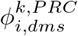 is also the actual location of the cell (i.e., 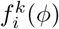 is maximal in response to radially facing light). Eq. 3 is plotted in Fig. 2A of the main text for PRCs in all five ambulacra (each ambulacrum in a different color).

The function 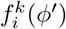 characterizes the response of PRCs to light coming from a punctiform source located at *ϕ*^′^ = *ϕ* −*ψ* (where *ϕ* is the location in the coordinate system of the arena). The response of the same PRC to a full stimulus comprising light coming from all directions *ϕ*^′^ with intensity *X*(*ϕ*^′^), was obtained by integrating the input *X* along the angular dimension, weighted by the angular sensitivity curve 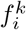:

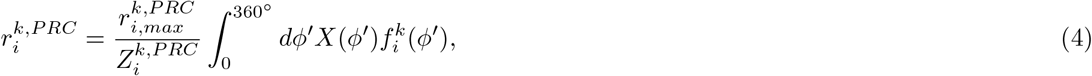

where 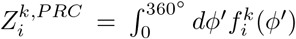 is a normalization factor to keep the output activity of PRCs in a physiological range (note that *X*(*ϕ*^′^≤) 1 for all stimuli; see the Supplementary Information, Sec. A.2, for mathematical details on the stimuli). The output is proportional to 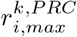, a constant parameter with physical units of PRC activity. Note that for our choice of stimuli and parameters, 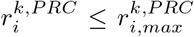. For an illustration of PRCs activity in response to stimuli, see the Supplementary Information, Sec. A.4.

#### 5.1.3 Radial nerve neurons

The activity of RNs is a sigmoidal function of the inhibitory input coming from the PRCs (through synapses *W*_*RP*_ *<* 0) and the excitatory input coming from adjacent RNs (through synapses *W*_*RR*_ *>* 0; see the main text). For convenience, we scaled the connection weights as the inverse of the square root of the number of projecting groups of cells:

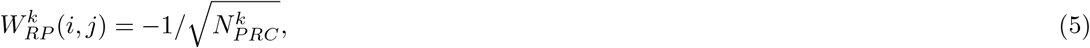

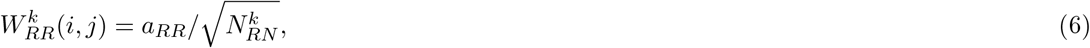

where *i, j* are the indices of the connected cells (with *i* being the index of the neuron on the receiving end of the connection), and the parameter *a*_*RR*_ quantifies the strength of ‘lateral’ excitation in the RN layer of each ambulacrum. The scaling with the inverse square root of the total number of neurons echoes wisdom from mammalian physiology (Barral and Reyes, 2016). In this model, however, this specific scaling is simply a means to parameterize the value of the connection strengths as a function of 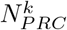 and 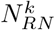.

The activity of RN neurons in response to input coming from both PRCs and other RN neurons, measured as firing rate (the number of action potentials per second), was

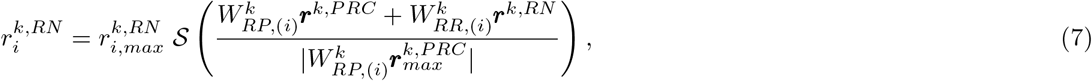

where 𝒮 (*x*) is the sigmoidal function Eq. 2 (Fig. 2B).

In Eq. 7, 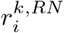 is the output firing rate of the *i*th RN cell on ambulacrum *k*, 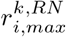 is the maximal firing rate (here, the same for all *i*), ***r***^*k,P RC*^ and ***r***^*k,RN*^ are vectors of activities from PRCs and RN cells, respectively, targeting RN cell *i* on ambulacrum *k* (these are visualized in Fig. 1B by the encircled groups of PRCs and RN neurons targeting the same group of RN neurons). Finally, we have used the notation *W*_(*i*)_ for the *i*th row of matrix *W*, and *W*_(*i*)_***r*** for the dot product of vectors *W*_(*i*)_ and vector ***r***: *W*(_*i*_)***r*** = ∑ _*j*_ *W*_*ij*_ *r*_*j*_.

#### 5.1.4 Oval nerve ring

Similar to RN neurons, eONR neurons received inhibitory and excitatory input (from adjacent iONR and eONR neurons, respectively; see the main text). Also in this case we rescaled the connection weights by the inverse square root of the total number of afferent neurons (note that 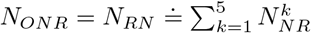)

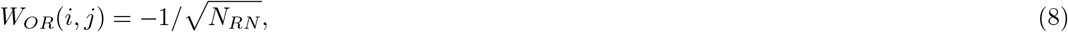

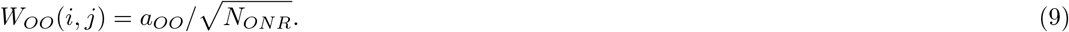

*W*_*OR*_ is the *effective*, inhibitory synaptic weight connecting RN neurons to eONR neurons (see Fig. 1B of the main text); *W*_*OO*_ is the lateral excitatory synaptic weight connecting adjacent groups of eONR neurons (Fig. 1B). Similarly to RN neurons, the output of eONR neurons is a sigmoidal function of their inputs,

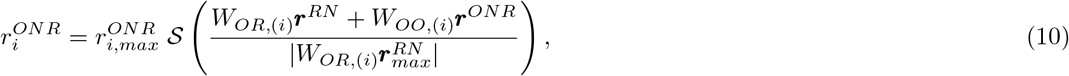

where 𝒮 (*x*) is Eq. 2 (parameter values given in Table 1 of the main text). In Eq. 10, the meaning of the symbols is analogous to the meaning of the corresponding symbols for RN neurons in Eq. 7. In particular, ***r***^*RN*^ and ***r***^*ONR*^ are vectors of activities from RN (on all ambulacra) and ONR cells, respectively, targeting (directly or indirectly) eONR cell *i*, according to the connectivity pattern shown in Fig. 1B of the main text.

#### 5.1.5 Population vector

The population vector of eONR cells was defined as the sum of vectors associated to each eONR neuron. The direction of each vector was the preferred direction of the corresponding neuron (defined below), while the length of the vector depended on the firing rate of the cell in response to the stimulus.

The preferred direction of eONR cell *i*, 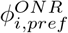, was defined as the direction of a narrow stimulus causing the maximal increase in activity in the cell. As the narrow stimulus, we chose a 2° white bar. The preferred direction can be identified with the unit vector 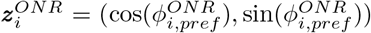, where we have used notation (*x, y*) for a vector with components *x* and *y*, respectively (note that 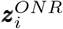 does not depend on the current visual stimulus: it depends only on the anatomy and physiology of the model). The population vector is the vector sum of the vectors 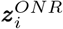, each weighted by the firing rate of the cell, 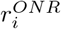:

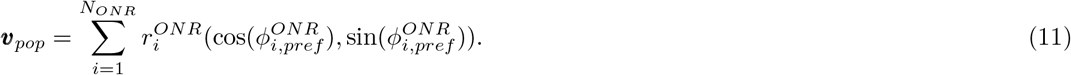

Unlike the single cells preferred directions, the population vector depends on the stimulus *X*(*ϕ*) (via the firing rates 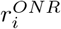) and on the orientation of the stimulus with respect to the animal (see Fig. 2CD of the main text).

Note that the preferred directions are properties of the cells and do not depend on the extended stimuli *X*(*ϕ*) used to probe movement; they can be computed analytically and agree with the preferred directions obtained in simulations with the 2° white bar (we report the analytical details in the Supplementary Information, Sec. A.3). The preferred direction of an eONR cell was highly correlated with its angular position in the coordinate system of the animal, although, due to the lack of PRCs in the region between ambulacra, a mismatch between eONR cell position and preferred direction slowly accrues until it is zeroed at the onset position of the next ambulacrum (see Supplementary Information, Fig. S2).

Each final position of the model on the wall of the arena shown in Fig. 4B was sampled from a circularly uniform distribution if the population vector was below threshold, and from a Gaussian distribution narrowly centered around the population vector if |***v***_*pop*_| *> θ*_*p*_ (with standard deviation 1*/*(|***v***_*pop*_| −*θ*_*p*_) ≤ 1; standard deviations 5 or 10 times larger gave the same results).

To compute the population vector in response to a visual stimulus we used Eq. 11 of the main text. Due to the recurrent connections in RNs and ONR, the neural activities need time to converge to the stationary values used in Eq. 11. Thus, we ran a simulation of the neural dynamics until the activity in the RN and ONR neurons reached the steady state. This process entails presenting the stimulus, obtaining the response of the PRCs, feed this response to the RN neurons, feed the RN neurons’ response to the iONR neurons, and so on until all neurons’ responses have been recorded. At the next step, the activity of the RN neurons is modified due to the lateral inputs coming from adjacent groups of RN neurons; and similarly for eONR neurons. The new activity in each group of neurons, in turn, modifies the activity of neighboring groups at the next time step, and so on. The cycle is repeated until a stable self-consistent activity state is found, i.e., a state where none of the neurons change their activity at the next time step. At this stage, we computed the population vector according to Eq. 11. We considered the steady state reached when the firing rates in two consecutive steps were smaller than 10^−5^. Note that convergence to steady state occurred rapidly: typically, in less than 200 iterations and never more than 600. Assuming a time step of 0.1ms for each iteration (a customary choice in the literature on neural systems), this means that convergence occurred typically in about 20ms and never more than 60ms, well below the typical time scales of locomotion in *D. africanum*.

#### 5.1.6 Model of movement

In brief, the model sea urchin will move in the direction of the population vector if the latter is sufficiently larger than the threshold *θ*_*p*_ = 5, otherwise it will move randomly but with a bias along the direction of the previous step. In detail, movement direction is a random variable that may follow one of two Gaussian distributions, *p*_1_(*μ*_1_, *σ*_1_) or *p*_2_(*μ*_2_, *σ*_2_), depending on the length of the population vector, *v*_*pop*_ = |***v***_*pop*_|. The distribution *p*_1_ is peaked around the direction of the population vector, while *p*_2_ is peaked around the previous direction of motion. In detail, *p*_1_(*μ*_1_, *σ*_1_) had a mean *μ*_1_ equal to the direction of ***v***_*pop*_ and 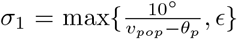, where *ϵ* = (10^−5^)° is a small number to keep the standard deviation *σ*_1_ positive. Also, values of *σ*_1_ larger than 360 were set to 360° (uniform distribution on the circle). For the Gaussian distribution *p*_2_(*μ*_2_, *σ*_2_), *μ*_2_ = *ϕ*_*pre*_ (the direction of motion at the previous step), with fixed standard deviation *σ*_2_ = 10° (on the first step, *ϕ*_*pre*_ was randomly sampled from a uniform distribution on the unit circle). The bias towards the previous direction helps to reduce the frequency of sharp directional changes, which are rarely observed experimentally.

The model followed *p*_1_ with probability

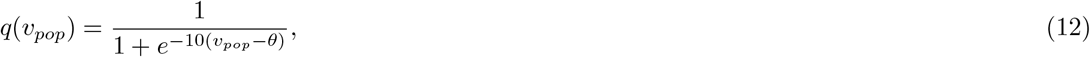

and it followed *p*_2_ with probability 1 −*q*(*v*_*pop*_), so that at every time step the movement is sampled from a mixture of two Gaussian distributions:

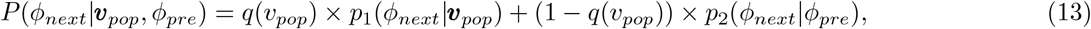

where *P* (*ϕ*_*next*_|***v***_*pop*_, *ϕ*_*pre*_) is the probability of the next step *ϕ*_*next*_ given the current value of the population vector (both direction and length) and the direction of the previous step, *ϕ*_*pre*_. For large population vector lengths *v*_*pop*_, *p*_1_ is almost certainly chosen, and it will be narrowly peaked around the direction of ***v***_*pop*_; on the other hand, if *v*_*pop*_ is below the threshold, the most likely movement will be a random step in a range of about 10 degrees from the previous direction. The probabilistic nature of this model is especially relevant for peri-threshold stimuli, for which one expects the largest behavioral variability since *p*_1,2_ have an equal probability of being selected.

To simulate the behavioral trajectories shown in Fig. 5 of the main text, each trial started with the animal located at the center of the arena with a random orientation with respect the center of the stimulus. The movement of the animal occurred in incremental steps covering 10% of the arena’s radius. The length of each step roughly mimics the number of steps required to reach the wall of the arena during the experiments. We also assumed the animal does not rotate during movement, as observed in experiment. After each step, we updated the position of the animal by sampling the new position from the distribution Eq. 13. Each trial ended when the animal reached the wall of the arena, i.e., when the center of the sea urchin reached a distance 3*/*4 from the center of the arena, as explained in the main text. Full details of the procedure, together with animations of behavioral trajectories, are given in Sec. A.5 of the Supplementary Information.

To obtain a dataset including *M* animals, we simulated *M* trials for each stimulus, each time with an animal starting from the center of the arena and with a uniformly random orientation of the first ambulacrum. As an aggregate measure of the final positions across a population of model sea urchins, we took the circular mean vector (Blevins and Johnsen, 2004; Kirwan et al., 2018)

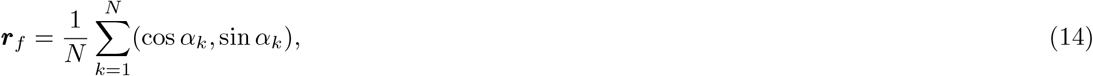

where *α*_*k*_ is the final position of model sea urchin #*k* (relative to the line joining the center of the area to the center of the stimulus), *N* is the total number of sea urchins, and we have used again notation (*x, y*) for a vector with components *x* and *y*, respectively. The circular mean vector ***r***_*f*_ is shown as the red vector in Fig. 4B of the main text. For a narrow concentration of final positions around a given direction, ***r***_*f*_ will point towards that direction and will have a large length (i.e., close to 1). Conversely, for a uniform distribution of final positions, ***r***_*f*_ will have a length close to zero. The Rayleigh test and V-test of circular statistics were conducted to test the significance of simulated object taxis at *α* = 0.05 confidence level (Figure 4B).

### 5.2 Statistical analysis

Model simulations and statistical analyses were performed in MATLAB using custom code. The Rayleigh test and V-test of circular statistics were conducted to test the significance of simulated object taxis (Figure 4B). The statistics were reported as mean values across 100 simulated experiments with 100 subjects in each experiment (where each subject is located at the center of the arena with a random initial orientation). Statistical results are in the caption of Figure 4B.

### 5.3 Code availability

The simulations and analysis for this study were performed with custom computer code written in MATLAB. The code is publicly available from https://github.com/lacameralab/diadema.

## A. Supplementary information

### A.1 Experimental setup of Kirwan et al. (2018)

Here we first briefly describe the setup for the taxis experiments reported in (Kirwan et al., 2018). Individual sea urchins of the species *D. africanum* were placed in a lit arena, surrounded by printed patterns containing a printed visual stimulus (Fig. 3). The arena comprised a cylinder of transparent acrylic and was surrounded by a white cylinder to exclude external cues. An array of four equidistant clusters of LEDs resulting in broad-spectrum visible illumination were placed above the arena. A remote-controlled camera was attached in an opening in an illumination diffuser at the top of the arena and was used to record time-lapse videos at a rate of 5 frames per second. The arena was filled with filtered natural seawater, at the same temperature at which the animals were housed (20°C). In each trial, the animal was placed by hand in the center of the arena and allowed to move to the periphery. Each trial continued for a maximum of 6 min or until the animal approached the arena wall. A trial was deemed complete if the animal moved at least three-quarters of the radial distance between the center and arena walls. Trials were conducted in sets of four and the stimulus was moved 90° clockwise for each subsequent trial, to remove the influence of any non-visual directional cues. Sets with individuals for which there were fewer than four completed trials (e.g. due to a loss of motivation) were excluded from analysis. The base of the arena was cleaned between trial sets with a brush to obscure chemical cues and the water was partially or completely changed, depending on its clarity. Experiments were performed during the daylight period of their entrainment. The frame rate for the recordings was 1 frame/s. Full details can be found in (Kirwan et al., 2018).

### A.2 Stimuli and coordinate systems

In the original experiments, the printed patterns surrounding the arena consisted of greyscale printed images, which were uniform in the vertical plane but in the horizontal plane included stimuli that consisted of dark regions set against a lighter background (Fig. 3B). The main patterns used a bar stimulus and a difference of Gaussians (DoG). These stimuli were used in the experiments of (Kirwan et al., 2018) analyzed here. The remaining stimuli were used in this paper to provide novel model predictions. In the bar stimulus, a region of homogeneously black stimulus was presented against a white background. In the DoG stimulus, the center of the stimulus was maximally dark, but of increasing reflectance towards the periphery of the stimulus (on the horizontal axis) and reaching the maximum achievable reflectance before darkening into the grey background. All stimuli were isoluminant with respect to the remainder of the patterns due to the lighter regions flanking the stimulus, i.e., it is not possible to detect the stimulus by simply comparing the radiance profile of different parts of the arena from the centre without having a spatial resolution equivalent to the arc subtended by the stimulus itself.

In the following we refer to the stimulus as the entire pattern surrounding the arena wall. The intensity of the stimuli along the longitudinal dimension (i.e., going from west to east on a horizontal plane) were indexed by an angle *ϕ* ∈ [0, 360)°, formed by an arbitrary reference line and the line connecting the center of the arena to the point of interest on the wall (we use degree (°) as the angle units in this paper). The normalized light intensity of a stimulus at *ϕ, X*_0_(*ϕ*) (henceforth simply ‘intensity’), indicates the reflectance of the stimulus at an angle *ϕ* from the center and varied linearly from 0.176 to 1 (Kirwan et al., 2018):

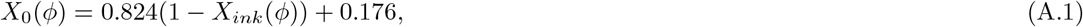

where *X*_*ink*_ was the ink value used in the actual experiments to print the images on the arena outer wall (Fig. 3B: the ink value of black is 1, for white is 0). Since photoreceptor cells respond to light, we replaced *X*_*ink*_ with 1− *X*_*ink*_ to convert the ink value to the intensity of light reflected by the stimulus in Eq. A.1. Note that in Eq. A.1 we have taken into account the relative reflectance of the paper (only a proportion of the light was reflected by the paper) (Kirwan et al., 2018). To obtain *X*_*ink*_ values between 0 and 1, we started from the functions *X*_*ink,raw*_ reported in Table S1, and then rescaled *X*_*ink,raw*_ by a simple linear normalization:

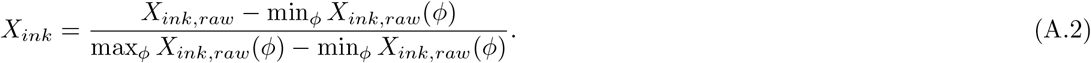

In addition to the stimuli above, we also used a homogeneous stimulus of constant intensity *X*_0_(*ϕ*) = 0.77 as control (same as the intensity of the DoG stimulus away from its center).

**Table S1:**
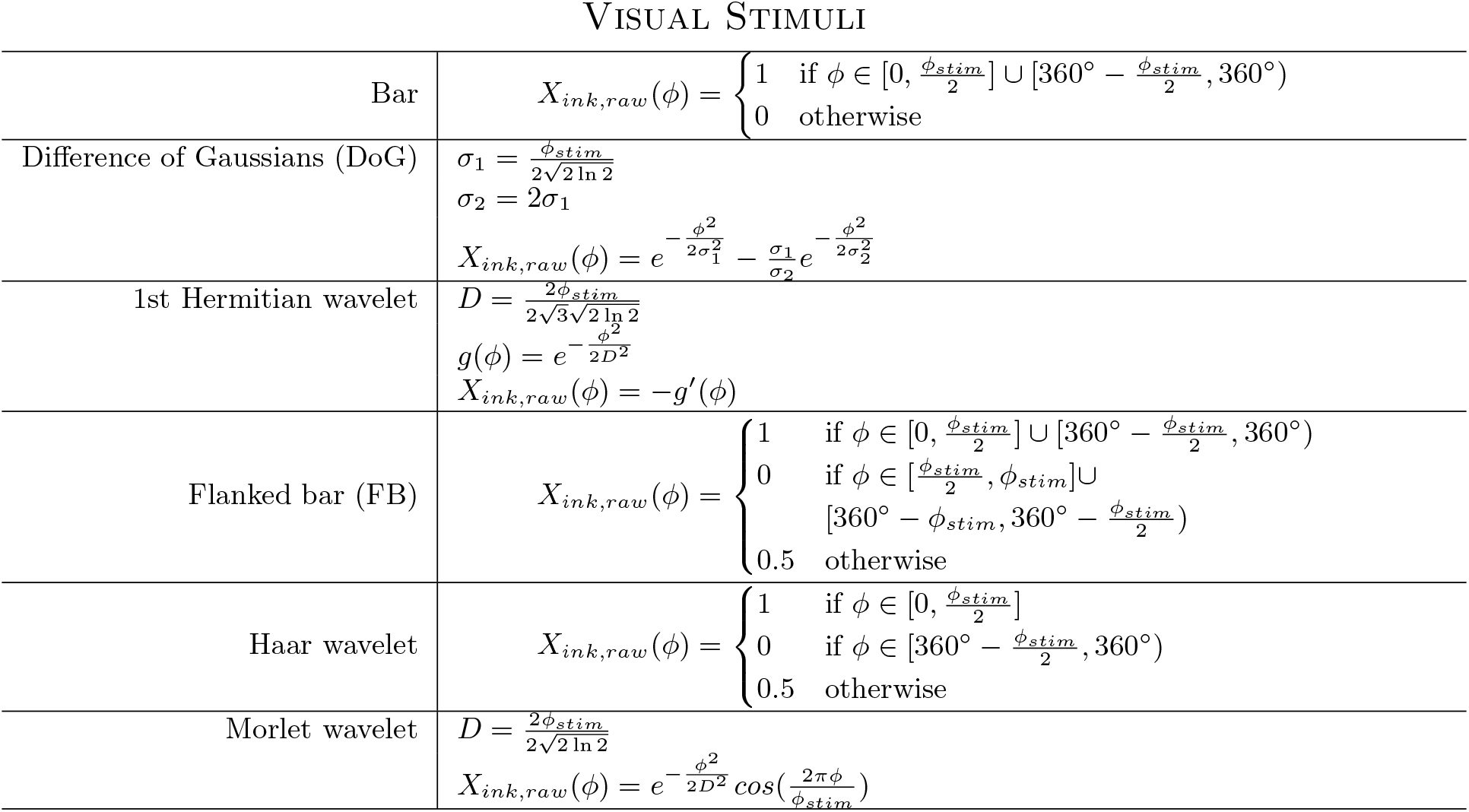
Mathematical definitions of the stimuli of Fig. 3B of the main text. The rightmost column contains the formulae for *X*_*ink,raw*_(*ϕ*) for each stimulus.

Note that each stimulus had an intuitively clear ‘center’ (Fig. 3B), which was always aligned to the reference line (i.e., the center of the stimulus was always at *ϕ* = 0° on the wall of the arena). The center of each stimulus was characterized by an arc width *ϕ*_*stim*_ which encloses the high amplitude region of the stimulus, surrounding the center. For example, in the case of the bar stimulus (see Fig. 3B), *ϕ*_*stim*_ corresponds to the arc width of the black region, while for the DoG it corresponded to the arc width of the distance between the two white maxima.

We next define the intensity of the stimulus in the animals’ own coordinate system, defined by the longitudinal angle *ψ* between the ‘first ambulacrum’ (arbitrarily chosen) and the origin of the arena’s coordinate system. The angle *ψ* defines the orientation of the animal in the arena, which is assumed constant during motion (i.e., the animal does not rotate as it moves; see Sec. A.5 for details). Therefore, each animal has its own (constant) value of *ψ*. This is illustrated in Fig. S1A, which shows a cartoon of the arena (black circle) with a sea urchin (purple circle) placed at the center of the arena. Black numbers outside the arena are coordinates in the arena’s coordinate system, while purple numbers are the ‘local’ coordinates in the sea urchin’s system. Given the local coordinate *ϕ*^′^ = *ϕ* −*ψ*, the intensity of light at angle *ϕ*^′^ is given by

**Figure S1:**
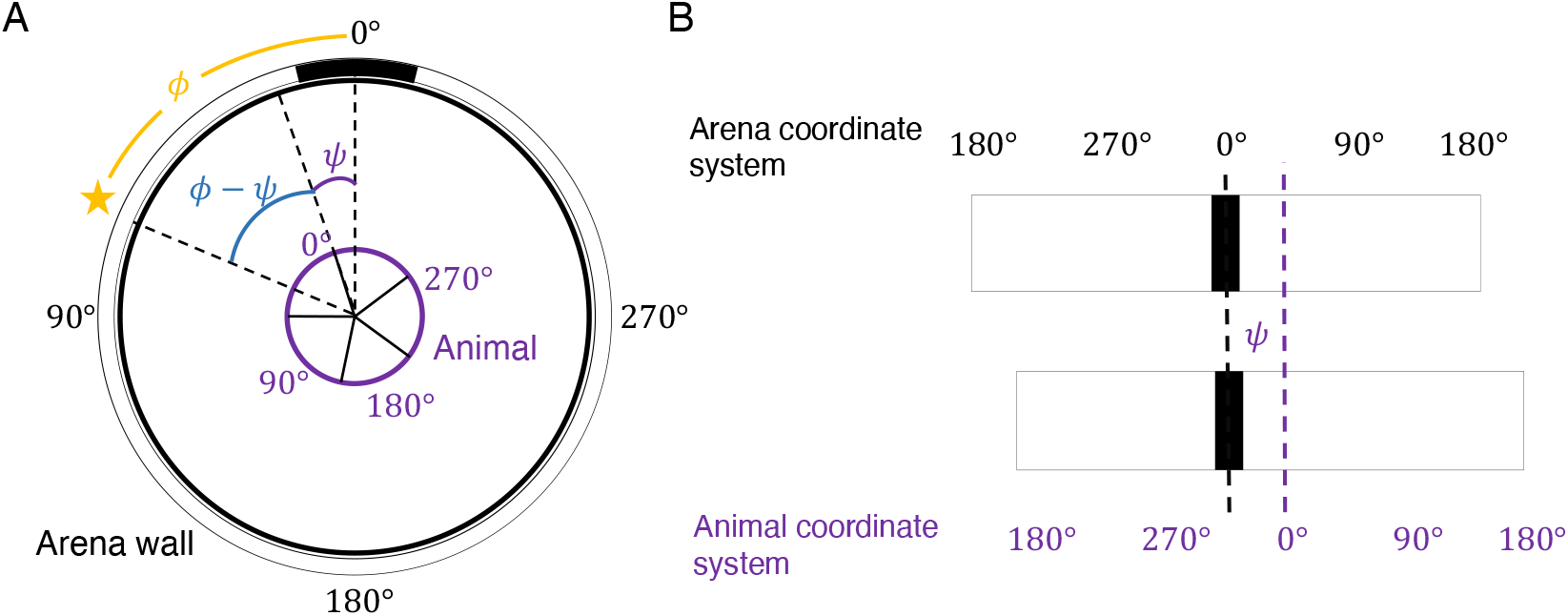
Light intensity in the reference frame of the sea urchin. **A**. Diagram of a *ϕ*_*stim*_ = 40° bar stimulus attached to the arena outer wall (black circle). Black numbers outside of the arena indicate the arena’s coordinate system. A sea urchin (purple circle) is placed at the center of the arena. Black solid lines originating from the center of the arena represent the ambulacra. One ambulacrum, here positioned at angle *ϕ* = 30°, is randomly chosen to be the ‘first ambulacrum’ and is the reference direction in the animal’s own coordinate system (purple numbers). The value of the ink at the location indicated by the yellow star is *X*_*ink*_(*ϕ*) = 0, and the light intensity at this location is *X*(*ϕ* − *ψ*) = *X*_0_(*ϕ*) = 0.824(1 − *X*_*ink*_(*ϕ*)) + 0.176 = 1. **B**. The top band is the stimulus paper in A cut at 180° in the arena’s coordinate system (black numbers at the top). The bottom band is the stimulus paper cut at 180° in the animal’s coordinate system (purple numbers at the bottom).

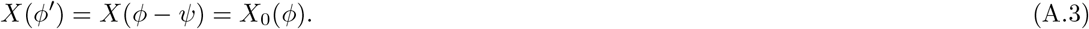

In our model, this quantity was the input to the photoreceptor cells distributed on the tube feet of the animal (Eq. 4 of Methods).

### A.3 Analytical formula for the preferred directions of eONR cells

Here we show how to compute analytically the preferred directions of eONR cells required in the definition of the the population vector, Eq. 11 of Methods. To do so, we associate unique vectors to each PRC, RN neuron and eONR neuron, and in each case define their preferred direction as the orientation of the associated vector. We then show that this definition matches the definition of preferred directions of eONR cells given in the main text.

First, consider the unit vector 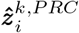 associated to PRC *i* on ambulacrum *k* having direction of maximum sensitivity 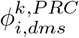 (see Sec. 5.1.2 of Methods):

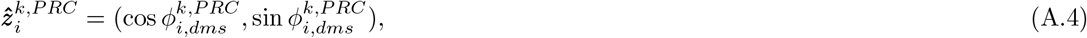

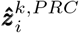 is a vector with unit length and orientation 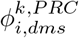.

We next associate a vector to each RN neuron by taking the linear combination of all PRC vectors connected to it, where the connection weights are used as coefficients of the linear combination:

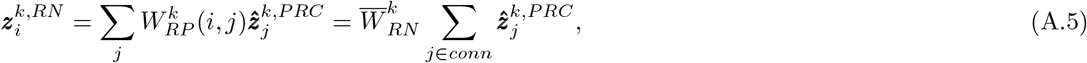

where *conn* is the set of PRCs connected to the RN cell *i* on ambulacrum *k*, and we have also used the fact that the non-zero connections *W*_*RP*_ (*i, j*) are all equal to the same value, here called 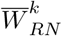 (see Eq. 5 of the main text). Since the lateral connections among RN neuron groups are symmetric with respect to the mean direction of maximum sensitivity of afferent PRCs, the RN neurons inherit a preferred direction (the orientation of vector 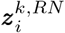) which is the same as the average direction of maximum sensitivity of its afferent PRCs.

We then associate a vector to each eONR cell in a similar way:

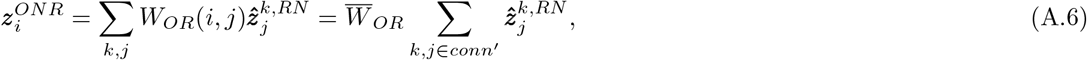

where 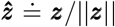, the sum over *k* is over the five ambulacra, and *conn*^′^ is the set of all RNs connected to the *i*th ONR cell via the ‘indirect pathway’ containing intermediate iONR cells (as shown in Fig. 1B of the main text). Again, the lateral connections among groups of eONR neurons do not impact the direction of 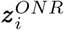, and therefore they are not included in the definition of 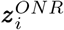.

By definition, the preferred direction of eONR cells is given by the orientation of their associated vectors 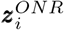. Fig. S2 shows that the preferred directions computed this way agree with the preferred directions obtained in response to a 2° white bar stimulus – the definition used in the main text.

Also note that 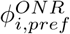 matches the orientation of the location of eONR cell *i* at the onset location of each ambulacrum (0°, 72°, 144°, …), and then a mismatch linearly accrues until the location of the next ambulacrum is reached. The mismatch is due to the lack of PRCs in the region between ambulacra, and its effect on the population vector is evident from the animations discussed in Sec. A.4.

We can now also write the population vector as a function of the PRC vectors. Recall that the population vector is defined as weighted sum of the eONR vectors (Eq. 11 of the main text), or, in the present notation,

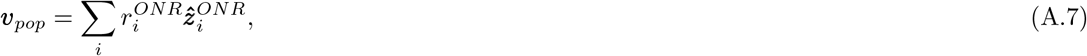

where 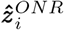 are the normalized vectors 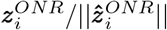. Putting together the steps leading to Eq. A.6, we obtain

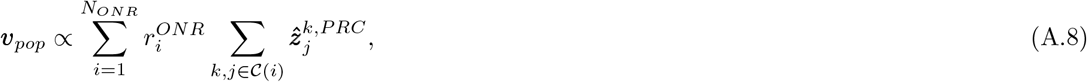

a vector sum of PRC vectors 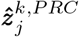 weighted by coefficients proportional to the firing rates 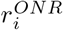. The sum over *k, j* ∈ C(*i*) is over all PRCs connected to eONR cell *i* via the ‘vertical pathway’ shown in Fig. 1B including intermediate RN and eONR cells. We note that this formula holds for our model but it does not necessarily hold for a different (especially non-symmetrical) topology of the neural connections from PRCs to ONR.

**Figure S2:**
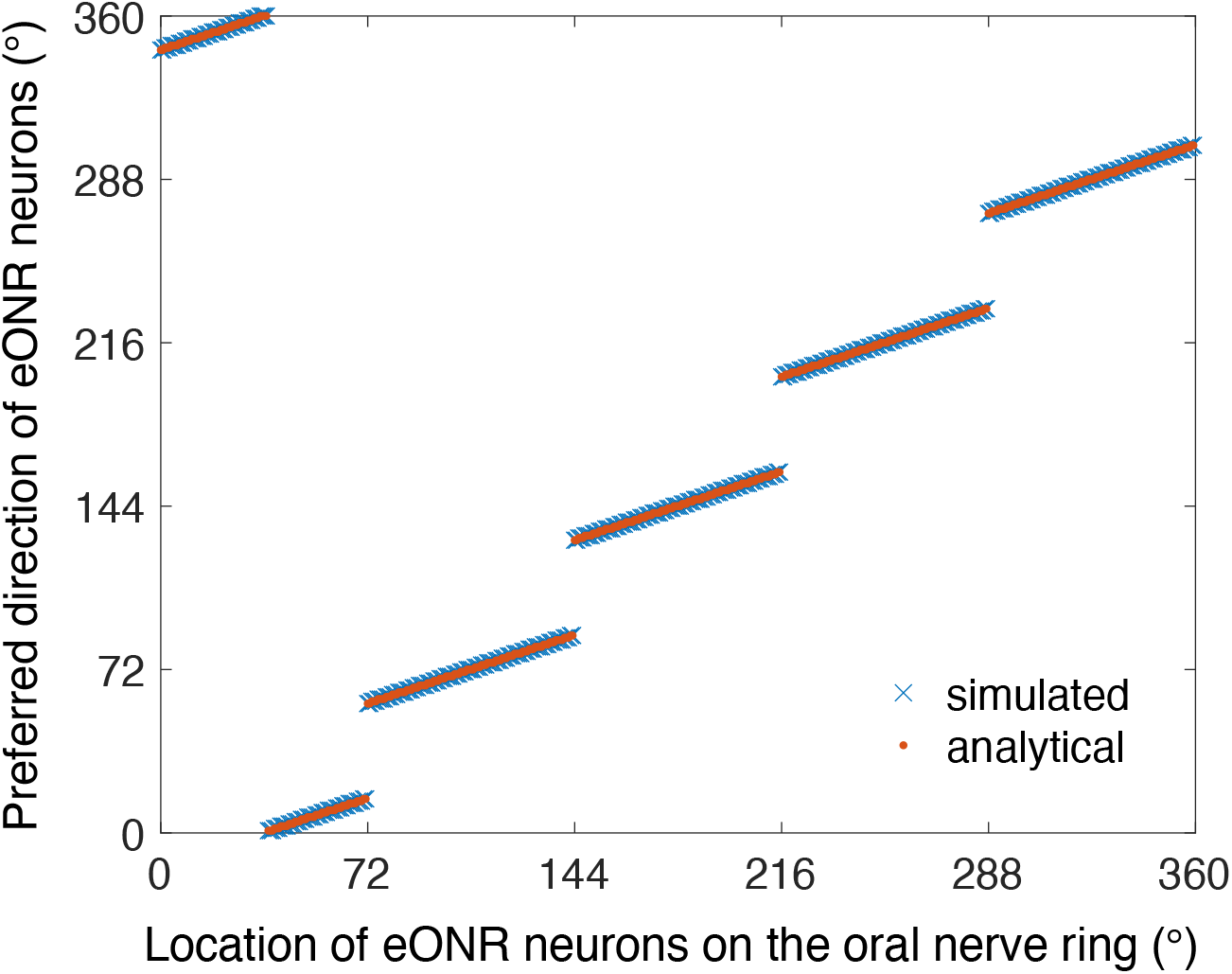
Comparison of preferred directions of eONR cells, defined as the orientation of vector A.6 (red circles) or as the direction of a 2° white bar stimulus that causes the maximal increase in activity in the eONR cell (blue crosses). The locations at 0°, 72°, 144°, 216° and 288° represent the locations of the ambulacra. Starting from each ambulacrum, the preferred direction is a linear function of the location of the eONR cell with slope smaller than 1. See the text for details.

### A.4 Neural activity and population vector as a function of the animal orientation

Table S2 lists the URL of three animations of the activity of PRCs, RN neurons, eONR neurons and the population vector as the relative position of the animal with respect to the stimulus changes. The stimuli used were the 40° bar, a 29° DoG and a 69° DoG. To better visualize this processs, we fixed the animal’s position and rotated the stimulus, which is equivalent to fixing the position of the stimulus and rotating the animal. Note how the orientation of the animal with respect to the stimulus affects stimulus detection due to the lack of PRCs between ambulacra. For narrow stimuli, this results in the existence of effective ‘blind spots’ between ambulacra.

**Table S2:**
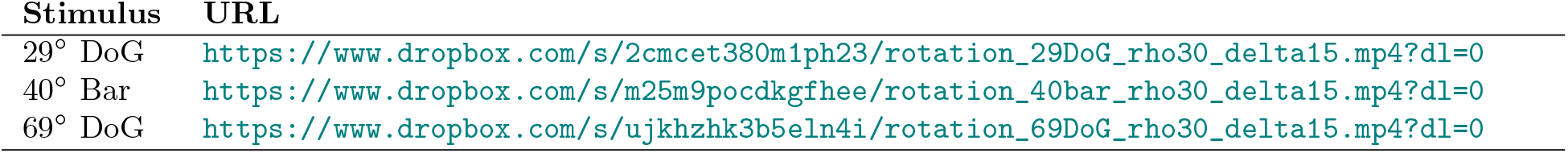
URLs of animations showing neural activity and population vector as a function of the relative orientation of the sea urchin and the center of the stimulus. Files also available from https://github.com/lacameralab/diadema/tree/main/diadema_animations.

### A.5 Simulation of behavioral trajectories

It is convenient to describe the model in the reference frame of the moving animal, where the location on the animal is described by an angle *ϕ* from its reference ‘first’ ambulacrum (see Sec. A.2). Each behavioral trial started with the animal located at the center of the arena with a random orientation *ψ* with respect the center of the stimulus (Fig. S3). Given a stimulus 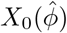 at location 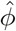 on the arena wall, the input to the PRCs was 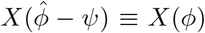 in the coordinate system of the sea urchin (this is Eq. A.3 after a change of notation 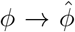 and *ϕ*^′^→ *ϕ*). We scaled the radius of the arena to be 1 and discretized the movement of the animal to occur in small steps *δr* = 0.1 at discrete points in time. Thus, rather than moving straight to the wall, the animal only made a small step and established the direction of the next step according to the stimulus detected in the current position. The length of each step roughly mimics the number of steps required to reach the wall of the arena during the experiments. We also assumed the animal does not rotate during movement as observed in experiment, in other words, the orientation of the unit vectors ***ê***_1_, ***ê***_2_ specifying the coordinate system of the animal (Fig. S3) does not change during locomotion, which empirically seems, at least approximately, correct. After each step, we updated the position of the animal by sampling the new position from the mixture of Gaussian distribution Eq. 13 of Methods.

The procedure is illustrated in Fig. S3. Starting at the center of the arena and facing stimulus *X*, the animal moves to a new position ***δr*** along, say, direction *α*: ***δr*** = (*δr* cos *α, δr* sin *α*). The intensity of the stimulus at the representative location ***x*** of the yellow star (in the coordinate system of the animal), is now detected at position ***x*** − ***δr***, forming an angle *ϕ*_*next*_ from the first ambulacrum. Given ***x*** = (cos *ϕ*, sin *ϕ*) and ***x*** − ***δr*** = (cos *ϕ* − *δr* cos *α*, sin *ϕ* − *δr* sin *α*) ≐ (*v*_*x*_, *v*_*y*_), the new angle *ϕ*_*next*_ is given by

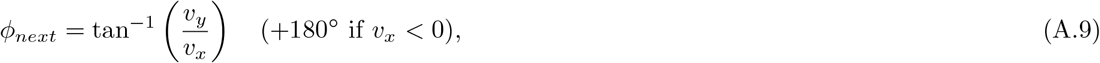

and the yellow star is detected as having intensity *X*_*next*_(*ϕ*_*next*_) from this new position. Note that in this new position, a different point on the wall faces the animal in the direction parallel to the ***x*** vector, that is, the stimulus appears as a new stimulus *X*_*next*_, which is then used to compute the next movement. The trial ended when the animal reached the wall of the arena. Since *D. africanum* has long spines, we modeled its radius as being 1*/*4 of the distance between the wall and the center of the arena. When the center of the sea urchin reached a distance 3*/*4 from the center of the arena, the animal had reached the wall and the simulation stopped. The current position at this point was projected orthogonally on the arena wall and logged as the final position.

Table S3 includes the URLs of three animations of the behavior of the model in response to a 40° bar, a 29° DoG and a 69° DoG. The urchin is represented by the moving circle with the 5 ambulacra drawn inside. The wall of the arena is represented by the large circle. The circle with the stimulus drawn on it and moving together with the urchin illustrates how the stimulus is ‘seen’ by the sea urchin at each new position. The red circle marks the threshold for the population vector (the latter always drawn in blue form the center of the arena). Each simulated trial ends when the edge of the urchin’s body (the smaller circle) makes contact with the arena.

**Figure S3:**
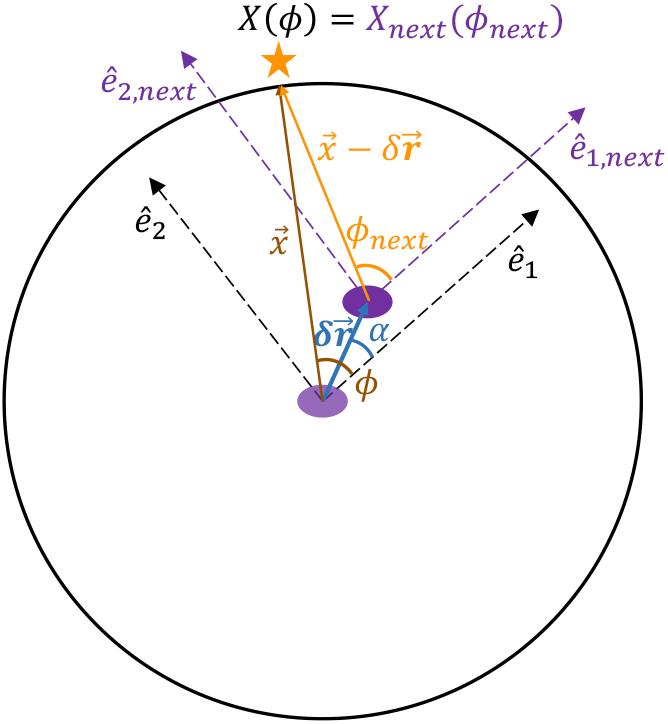
Stimulus coordinates during simulation of movement. The unit vectors ***e***_1_ and ***e***_2_ are the principal axes of the coordinate system of the sea urchin (purple ellipsoid). ***ê***_*i*_ is the direction of the first ambulacrum in the coordinate system of the arena. The animal moves without rotating, so that the orientation of its axes remains constant (i.e., ***ê***_1,*next*_ = ***ê***_1_ and ***ê***_2,*next*_ = ***ê***_2_). The relative position of the yellow star with respect to the sea urchin is at *ϕ* when the animal is at the center of the arena, and at position *ϕ*_*next*_ after the animal has made one step ***δr***. From the new position, the stimulus appears as a new stimulus *X*_*next*_, which is then used to compute the next movement, and so on until the animal reaches the wall of the arena and its final position is recorded.

**Table S3:** URLs of animations showing simulations of the behavior of the model in the presence of a 40° bar, a 29° DoG and a 69° DoG. All tracks comprising 100 simulations per stimulus are shown in Fig. 5 of the main text. Files also available from https://github.com/lacameralab/diadema/tree/main/diadema_animations.

| Stimulus | URL |
| --- | --- |
| 29° DoG | https://www.dropbox.com/s/x9jbp04g0hpkjx7/trajectory_29DoG_N10_rho30_delta15.mp4?dl=0 |
| 40° Bar | https://www.dropbox.com/s/eh1jlk3n9fpzkez/trajectory_40bar_N10_rho30_delta15.mp4?dl=0 |
| 69° DoG | https://www.dropbox.com/s/fylhsr1a7oeaayf/trajectory_69DoG_N10_rho30_delta15.mp4?dl=0 |

### A.6 The effect of variability in PRCs’ acceptance angles

The results of the main text (summarized in Fig. 6) were obtained using a constant acceptance angle for PRCs. Here we repeat the analysis of section “Effect of location and acceptance angle of PRCs on spatial vision” of the main text and show that our results also hold in the case of random distributions of acceptance angles across PRCs. For concreteness, we chose, in each case, Gaussian distributions with mean ⟨Δ*ρ*⟩ and standard deviation equal to ⟨Δ*ρ*⟩ */*4 (different choices did not qualitatively alter the results). The results are shown in Fig. S4. Except for noisier contours, the regions of parameter space defined by the *v*_*max*_ ≥ 4 or *v*_*max*_ ≥ 5 appear unchanged compared to those shown in Fig. 6 of the main text for constant Δ*ρ*.

**Figure S4:**
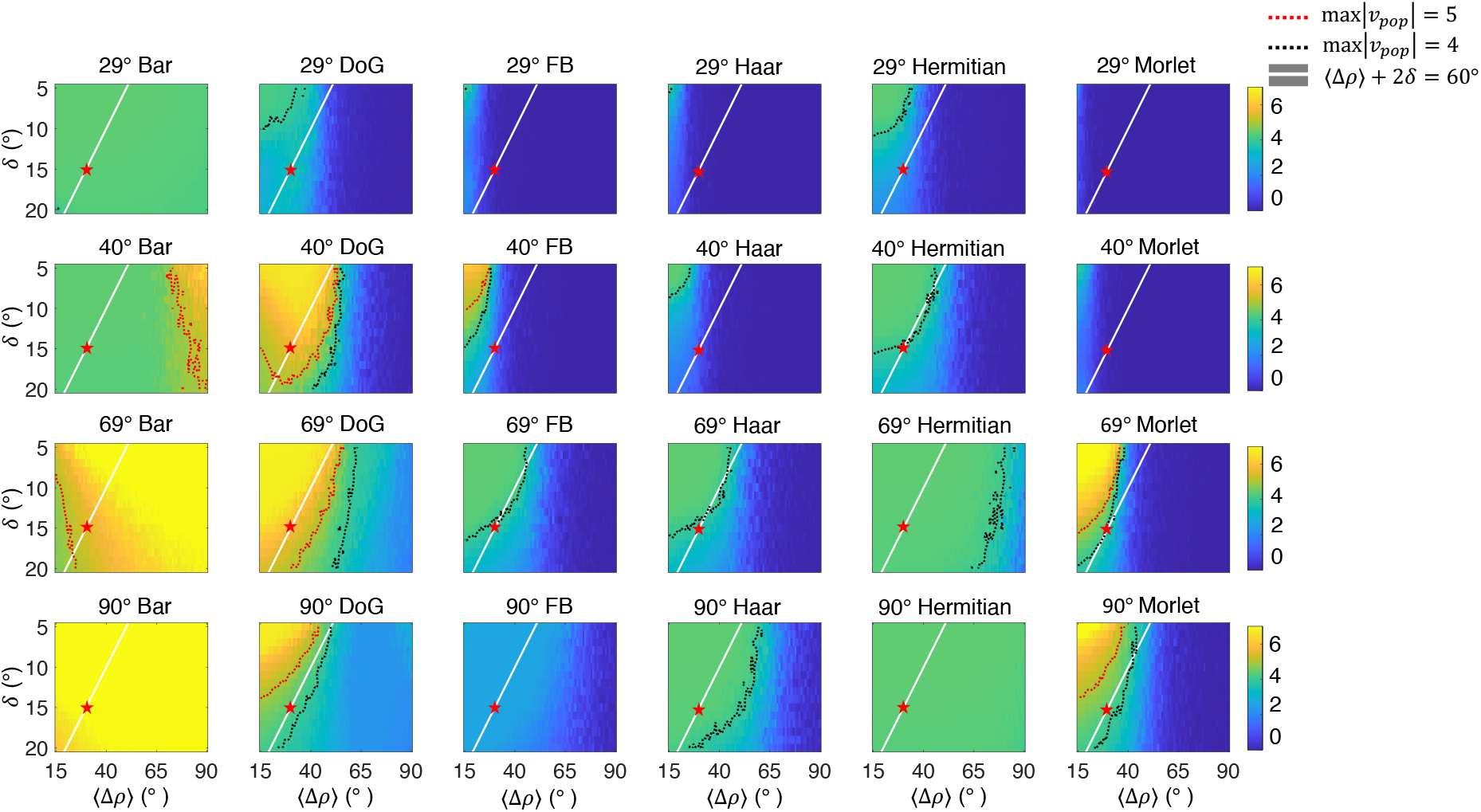
Effect of acceptance angle and location of PRCs on the model’s spatial vision. Same as Fig. 6 of the main text but with a random distribution of PRC acceptance angles Δ*ρ*, chosen to be a Gaussian distribution with mean ⟨Δ*ρ*⟩ and standard deviation ⟨Δ*ρ*⟩*/*4. Each panel shows a heat map of *v*_*max*_, the maximal length of the population vector across initial orientations of the animal, for a given stimulus and a given pair of values for ⟨Δ*ρ*⟩ and *δ*. Each column shows the same stimulus for different arc widths of the target region (i.e., for different *ϕ*_*stim*_, see Table S1), while each row shows the same *ϕ*_*stim*_ across different stimuli. If *v*_*max*_ < *θ*_*p*_, the animal cannot detect the stimulus from any orientation. The red dotted line is the contour line where *v*_*max*_ = *θ*_*p*_, while the black dotted line is the contour line where *v*_*max*_ = 4. The white line is the collection of points with ⟨Δ*ρ*⟩+2*δ* = 60°. Red stars mark the point (⟨Δ*ρ*⟩, *δ*) = (30°, 15°), the parameter values used in the main simulations (where, however, Δ*ρ* was constant across PRCs). *DoG*: Different of Gaussians; *FB*: Flanked bar.

